# Functional divergence of regulatory and conventional bovine neutrophils following *Mycobacterium bovis* infection

**DOI:** 10.64898/2026.03.16.712159

**Authors:** Maïa Saint-Vanne, Badreddine Bounab, Sebastien Eymieux, Emma Perdriau, Florence Carerras, Romain Roullier, Yves Le Vern, Julien Pichon, Emilie Doz-Deblauwe, Pierre Germon, Nathalie Winter, Aude Remot

**Author notes:** Corresponding Author: Aude Remot.

## Abstract

Bovine tuberculosis (bTB), due to *Mycobacterium bovis* (Mb) infection, is a chronic cattle disease and neglected cause of zoonotic tuberculosis. The role of neutrophils in bTB is overlooked. We recently identified a new neutrophil subset in cattle and mice with a similar morphology to conventional inflammatory neutrophils (N^conv^). However, unlike N^conv^, these regulatory neutrophils (N^reg^) express MHC-II at their surface and can suppress lymphocyte proliferation. In this study, we compared the responses of bovine N^conv^ and N^reg^ to infection with a virulent Mb strain circulating in France and the attenuated Mb BCG vaccine. N^reg^ and N^conv^ had different transcriptional profiles and were differentially activated by Mb infection. Both N^reg^ and N^conv^ efficiently killed Mb, but N^reg^ had higher levels of phagocytosis activity and ROS production. N^reg^ had higher levels of mitochondrial activity and an ultrastructural organization different from that of N^conv^. Our results provide the first insight into the functional characterization of bovine neutrophil subsets during Mb infection and highlight a new layer of complexity in their functional diversity that must be taken into account to improve our understanding of bTB pathophysiology, which is urgently required to improve the management of this costly disease.

## Introduction

Bovine tuberculosis (bTB), due to *Mycobacterium bovis* (Mb) infection, is a zoonotic disease and one of the most difficult infections to control in cattle. This disease is a major public health issue with a large global economic burden in both developing and developed countries, due to the lack of antituberculosis vaccines for livestock [1], and the lack of reliability of diagnostic techniques [2,3]. bTB is a notifiable disease dealt with by the culling of entire herds [4], resulting in a large economic, social, ethical and environmental impact. France has been officially bTB-free since 2001, with a prevalence of less than 0.1% in cattle herds [5,6], but this status is threatened by the many Mb strains circulating among wildlife and livestock in some parts of the country.

The pathophysiology of bTB has been described in less detail than that of human TB, caused principally by *Mycobacterium tuberculosis* (Mtb). The correlates of protection remain unknown for both human and animal TB, and their identification is a matter of top priority for the World Health Organization and the World Organization for Animal Health. Improvements in our understanding and control of the disease require a deeper understanding of the immune mechanisms involved during infections in cattle. The role of macrophages in the pathophysiology of this disease is well established [7–9], but that of neutrophils — the other main type of phagocytic immune cell — remains largely unknown. However, neutrophils have been shown to play important roles in innate immune resistance to Mtb infection in both humans and mouse models [10]. Indeed, individuals in contact with patients presenting active TB are less likely to be infected with Mtb if they have high peripheral blood neutrophil counts [10]. Conversely, neutrophils constitute the largest cell population in the lungs of patients with active TB [11] and the transcriptional signature of severe TB is driven principally by type I IFN signaling, dominated by neutrophils [12]. This neutrophil-driven IFN signature is also validated at the protein level [13]. In Mtb-resistant mouse models, two different waves of neutrophils are observed after Mtb infection, before and after the onset of adaptive immunity [14]. The neutrophils of the first wave take up mycobacteria by phagocytosis *in situ* in the lung, whereas the second wave is T cell-dependent, with neutrophils seldom associating with the bacilli, instead establishing close contact with T cells [14], helping to form the structured mature granuloma [15,16]. Early neutrophils generated during the innate phase of the response to infection are therefore key to the containment of the bacilli, either because they kill the bacilli directly, or because they contribute to the formation of the beneficial granuloma that restricts bacterial multiplication. By contrast, uncontrolled and chronic neutrophil influx is also associated with an inflammation burst, tissue damage and a poor clinical outcome in active TB [17,18].

Neutrophils were long considered to be homogeneous, mostly pro-inflammatory cells capable of reaching sites of infection and rapidly eliminating microorganisms and other dangers. However, this restricted view of neutrophils has evolved considerably over the last decade. It is now well established that neutrophils are heterogeneous [19–21] and form discrete subtypes with a broad range of functions, including anti-inflammatory [22,23] and regulatory roles [24]. We recently identified a new subset of neutrophils — “regulatory” (N^reg^) neutrophils — circulating in both cattle and the mouse model [25]. Like conventional pro-inflammatory neutrophils (N^conv^), N^reg^ have a classical polymorphonucleus and these two types of neutrophils have similar cell-surface markers. However, unlike N^conv^, N^reg^ express major histocompatibility complex class II (MHC-II) on their surface, together with PD-L1 in mice. At steady state, N^reg^ can suppress T lymphocyte proliferation, thereby acting as key partners in the adaptive immune response. We show that N^conv^ exacerbate local inflammation in mouse models of TB, through caspase-1-dependent IL-1β production, accelerating inflammation by maintaining a vicious circle of inflammatory neutrophils and CD4^+^ T cells. By contrast, N^reg^ act through PD-L1 to block T cell proliferation and IFN-γ production, thereby reducing inflammation [26].

Our knowledge of N^reg^ function during infections in cattle remains limited. We recently demonstrated that both the N^reg^ and N^conv^ subsets were present in milk from healthy Holstein and Normande cows and that, during mastitis, milk N^reg^ had a greater bactericidal capacity than their classical counterparts [27]. We also observed a positive correlation between the counts of N^reg^ and T cells in milk during infection.

Neutrophils play crucial roles in the pathophysiology of TB that differ according to disease stage [28]. We therefore further investigated the functions of the newly described N^reg^ in controlling Mb at early stages of disease. We used two strains of *M. bovis* — the virulent Mb3601 strain circulating in France and the attenuated BCG vaccine strain — to confirm that N^conv^ and N^reg^ had different transcriptional signatures. We identified key biological pathways that differed markedly between the two neutrophil subsets. We then compared the functions of N^reg^ and N^conv^, including bacterial killing, reactive oxygen species (ROS) production and phagocytosis. Finally, we performed electron microscopy to compare the ultrastructure of the two subsets of neutrophils. These analyses suggested that the two neutrophil subsets interact differently with Mb *in vitro.* Our data pave the way for a reconsideration of the role of neutrophils in the pathophysiology of bTB.

## Materials and Methods

### Bacterial Strains – Growth Conditions and Inoculum Preparation in a Biosafety Level 3 Laboratory

Mb3601 is a French Mb strain isolated by the French bTB reference laboratory at ANSES from an infected cow in Bourgogne-Franche-Comté. The BCG-WT 1173P2 vaccine strain was obtained from the Pasteur Institute. Recombinant fluorescent BCG mCherry and Mb3601 mCherry strains were obtained by electroporation with the pNIP40-mCherry integrative plasmid and selection on hygromycin B (50 μg/mL, Sigma), as previously described [29]. Bacteria were grown in flasks containing 7H9 broth, 10% ADC, 0.05% Tween 80, and sodium pyruvate (38 mM). At mid-exponential growth phase, the bacteria were harvested, tittered, dispensed into aliquots and stored at - 80°C until use. BCG-WT batch titers were determined by plating serial dilutions on 7H11 agar supplemented with 0.5% glycerol and 10% OADC (ADC supplemented with 0.005% oleic acid). Mb3601 batch titers were determined by plating serial dilutions on 7H11 agar supplemented with 10% OADC, 10% calf serum and 0.005% decomplemented sheep’s blood diluted 1:2 in sterile milliQ water. The plates were incubated at 37°C for up to 3 weeks and the colony-forming units (CFUs) were then counted. For each strain and each experiment, inoculum was prepared by thawing one frozen aliquot in 7H9 medium, 10% ADC, 0.05% Tween 80 and incubating it for half a day at 37°C. The aliquot was then centrifuged for 10 min at 1,700 x *g*, and bacterial concentration was adjusted with RPMI medium, as appropriate, for each experiment.

### Cow’s blood samples

Blood samples were collected from primiparous Prim’Holstein cows from the Orfrasière Animal Physiology Experimental Unit (UE-PAO) at INRAE Centre Val-de-Loire. Blood samples were collected from the jugular vein into 10 mL K2 EDTA Vacutainer tubes and were kept at room temperature until processing (<1 hour). All the cows were housed together and our protocol did not interfere with routine herd management. Blood sampling was approved by the Ethics Committee for Animal Experimentation Val-de-Loire (CEEA VdL), registered with the French National Committee for Animal Experimentation (APAFIS no. 202007241825481.V2), and complied with EU regulation 2010/63.

### Cell Preparation from Blood Samples

Tubes containing blood were centrifuged at 1000 x *g* for 30 min at 20°C (acceleration 9/brake 7). The plasma layer and the buffy coat were removed. ACK lysis buffer (Thermo Fisher Scientific, 4 vol buffer/1 vol of blood) was added and the mixture was incubated for 5 minutes at room temperature to lyse the red blood cells. The remaining cells were washed twice with Dulbecco phosphate-buffered saline (D-PBS) and then resuspended in FACS buffer (D-PBS, 2 mM EDTA, 10% horse serum) for further labeling, or neutroRPMI (RPMI supplemented with 2 mM L-glutamine, 10 mM HEPES, and 1 mg/mL BSA with extremely low endotoxin level (≤2.5 endotoxin units/mL, Sigma) for cell culture. Cells were stained with Acridine Orange fluorescent stain (Logos Biosystems) and counted with a Luna-FL^TM^ Automated Fluorescence Cell Counter (Logos Biosystems).

### Cell Sorting by Flow Cytometry

All the antibodies and probes used in this study are listed in **Table 1**, together with the clones, suppliers, and working concentrations or dilutions. Blood cells were labeled by incubation with the primary antibodies (G1 and MHC-II) for 20 min at 4°C. Cells were washed in D-PBS, labeled with the corresponding fluorescently labeled secondary antibody (conjugated with Alexa Fluor 647 and Alexa Fluor 488) and a Fixable Viability Dye (FVD-eFluor 780). Cells were then resuspended in neutroRPMI and live neutrophils (FVD^-^ G1^+^ MHC-II^-^ and FVD^-^ G1^+^ MHC-II^+^ cells) were obtained by sorting on a MoFlo Astrios EQ high-speed cell sorter (Beckman Coulter) according to the protocol described by Rambault et al (2021) [30].

**Table 1:** List of antibodies and probes used in this study.

| Target or name | Isotype | Clone | Working concentration or dilution | Reference | Supplier |
| --- | --- | --- | --- | --- | --- |
| Anti-bovine G1 | Mouse IgM | CH138A | 2 µg/mL | WSC0608B-100 | Clinisciences |
| Anti-feline MHC-II | Mouse IgG1 | CAT82A | 2 µg/mL | WSC0525F-100 | Clinisciences |
| Anti-mouse IgM Alexa Fluor 647 | Goat IgG | Polyclonal | 10 µg/mL | A21238 | Thermo Fisher Scientific |
| Anti-mouse IgG1 Alexa Fluor 488 | Goat IgG | Polyclonal | 10 µg/mL | A21121 | Thermo Fisher Scientific |
| Anti-mouse IgG1 phycoerythrin (PE) | Rat IgG | RMG1-1 | 1 µg/mL | 406608 | BioLegend |
| Fixable Viability Dye (FVD) eFluor 780 | - | - | 1:10,000 | 65-0865-14 | Thermo Fisher Scientific |
| Viability dye Zombie Aqua | - | - | 1:500 | 423102 | BioLegend |
| Sulforhodamine-FLICA Polycaspase | - | - | 1:250 | ICT916 | Bio-Rad |
| CellROX Green | - | - | 5 µM | C10444 | Thermo Fisher Scientific |
| MitoTracker Orange | - | - | 250 nM | MT7510 | Thermo Fisher Scientific |
| Hoechst 33342 | - | - | 20 µM | 62249 | Thermo Fisher Scientific |

### RNA Extraction

Sorted neutrophils were infected with BCG (BCG-WT at a multiplicity of infection (MOI) of 10) and Mb (Mb3601 at a MOI of 1) strains. The plates were incubated for 2 h at 37°C. The supernatant was then discarded and Tri-Reagent was added to each well. Technical duplicates or triplicates were pooled in an Eppendorf tube and kept at-80°C until RNA extraction. The initial step of RNA extraction consisted of a pre-extraction with chloroform (5:1, vol/vol), followed by vigorous shaking and incubation for 10 min at room temperature. The samples were then centrifuged at 12,000 x *g* for 10 min at 4°C, and the aqueous phase was collected and mixed with isopropanol (1:1, vol/vol). Total RNA was then purified using the NucleoSpin RNA kit (Macherey-Nagel) in accordance with the manufacturer’s instructions, with a DNase treatment step.

### RNA Sequencing and Differential Expression Analysis

RNA sequencing was performed by an external service provider (Helixio, France, https://www.helixio.fr/page/rna-seq). RNA integrity was assessed before sequencing (RIN ranging from 5.7 to 9.6 (mean ± SEM: 7.70 ± 0.22)). Raw paired-end reads were assessed with FastQC (v0.12.1) [31]. Adapter sequences and low-quality bases were trimmed off with fastp (v0.23.4) [32] in paired-end mode, and only cleaned paired reads were retained. Post-trimming read quality was evaluated with FastQC. Transcripts were quantified by pseudoalignment with the *Bos taurus* transcriptome (ARS-UCD1.3 assembly, Ensembl cDNA release 113) in Salmon (v1.10.0) [33], using an index built with a k-mer size of 31. The --validateMappings option was enabled, and bias corrections were applied for effective transcript length, sequence composition, and GC content [33]. Transcript-level abundance estimates (quant.sf files) were imported into R (v4.4.2) with tximport (v1.34.0) [34] and summarized at gene level with a transcript-to-gene mapping table derived from the Ensembl annotation, ignoring transcript version suffixes. The resulting gene-level counts were used to construct *DESeqDataSet* objects with DESeq2 (v1.46.0) [35]. Weakly expressed genes (fewer than 5 normalized counts in at least 6 samples) were filtered out before differential expression analysis. Size factors were estimated with DESeq2, and variance-stabilizing transformation (VST) was applied for exploratory analyses. Principal component analysis (PCA) was performed to visualize the global structure of the dataset by neutrophil subtype (N^conv^ *vs* N^reg^) and experimental conditions. Differential expression between regulatory (N^reg^) and conventional (N^conv^) neutrophils was assessed separately for each set of experimental conditions (uninfected, BCG, Mb3601). For each set of experimental conditions, a DESeq2 model with the design formula ∼CellType was fitted, using N^conv^ as the reference. Log_2_ fold-changes were moderated by the *apeglm* method [36]. Genes were considered differentially expressed if the adjusted *p*-value (Benjamini–Hochberg method) was < 0.05 and |log_2_FC| ≥ 0.58 [37].

Functional enrichment analyses were then performed separately for each gene set derived from condition-specific and intersecting signatures, distinguishing genes upregulated in N^reg^ and N^conv^. Enrichment analyses were performed with g:Profiler (gprofiler2 package [38,39]) for *Bos taurus*, applying an overrepresentation analysis (ORA) framework and querying the Gene Ontology (Biological Process, Molecular Function, Cellular Component [40,41]), Reactome and KEGG databases. False-discovery rate correction was applied, and terms displaying significant enrichment were visualized on dot plots.

The transcriptomic data have been deposited in the European Nucleotide Archive (ENA) under BioProject accession number PRJEB106984.

### Cell Survival

Blood cells were labeled with primary antibodies (G1 and MHC-II, see **Table 1**) for 20 min at 4°C. Cells were washed in D-PBS and labeled with the corresponding fluorescent secondary antibody (conjugated with Alexa Fluor 647 or Alexa Fluor 488) and the Zombie Aqua viability dye. Cells were washed in D-PBS and resuspended in neutroRPMI. Neutrophil apoptosis was evaluated with the Sulforhodamine (SR-)FLICA^TM^ Poly Caspase Kit according to the manufacturer’s instructions. Briefly, cells were infected with BCG-WT (at a MOI of 10) or Mb3601 (at a MOI of 1), with the addition of SR-FLICA^TM^ Poly Caspase reagent at the same time as the bacteria. Staurosporine (STR, working concentration 2 µM) was used as a positive control for this experiment. The plates were incubated at 37°C for various times: 30 min, 2 h, 4 h and overnight. The cells were then fixed by incubation with 4% PFA for at least 2 h. Fluorescence was measured with an LSR Fortessa^TM^ X-20 Flow Cytometer (BD Biosciences). Flow cytometry data were analyzed with Kaluza Analysis software (2.1 version, Beckman Coulter).

### Killing Assay

Sorted neutrophils were infected with Mb (BCG-WT at a MOI of 10 or Mb3601 at a MOI of 1). The plates were incubated for 2 h at 37°C. The supernatant was collected and the cells were lysed in D-PBS supplemented with 0.1% Triton X-100. Both supernatant and lysate were plated at appropriate dilutions on plates containing solid medium of the composition described above. Killing was assessed 2 h after infection, by counting CFUs in the supernatant and cell lysate. The percent killing was determined as CFUs in the supernatant + CFUs in the neutrophil lysate, multiplied by 100 and divided by the total number of bacteria in the inoculum, which was verified for each experiment.

### Phagocytosis Assay

Blood cells were labeled by incubation with primary antibodies (G1 and MHC-II) for 20 min at 4°C. The cells were washed in D-PBS and labeled with the corresponding fluorescent secondary antibody (conjugated with Alexa Fluor 647 and Alexa Fluor 488) and the FVD-eFluor 780. The cells were washed in D-PBS and resuspended in neutroRPMI. Labeled cells were then infected with recombinant BCG-mCherry (at a MOI of 10) or recombinant Mb3601-mCherry (at a MOI of 1). At various time points, infected cells were centrifuged at 400 x *g* for 5 min and the supernatant was discarded. We then added 4% paraformaldehyde (PFA) to each tube and incubated the tubes overnight. Fluorescence was measured with a LSR Fortessa^TM^ X-20 Flow Cytometer. Flow cytometry data were analyzed with Kaluza Analysis software, 2.1 version.

### ROS Production Assay

Blood cells were labeled by incubation with primary antibodies (G1 and MHC-II) for 20 min at 4°C. in the cells were washed in D-PBS and labeled with the corresponding fluorescent secondary antibody (conjugated with Alexa Fluor 647 and phycoerythrin) and the FVD-eFluor 780 viability dye was added. The cells were washed with D-PBS and resuspended in neutroRPMI. ROS production by neutrophils was assessed with the CellROX^TM^ Green Flow Cytometry Assay Kit according to the manufacturer’s instructions. Briefly, cells were infected with BCG-WT (at a MOI of 10) or Mb3601 (at a MOI of 1), with the addition of CellROX Green^TM^ at the same time as the bacteria. The plate was incubated for 2 h at 37°C. The cells were then fixed by incubation with 1% PFA overnight. ROS fluorescence was measured with an LSR Fortessa^TM^ X-20 Flow Cytometer. Flow cytometry data were analyzed with Kaluza Analysis software, 2.1 version.

### Mitochondrial imaging by confocal microscopy and image analysis

Sorted neutrophils were placed in an 18 well microslide (IBIDI) and incubated with MitoTracker Orange CMTMRos for 30 min and Hoechst 33342 for 10 min at 37°C. The cells were then washed with D-PBS and imaged by confocal laser scanning microscopy (CLSM) (AX NSPARC with WI 60x/1.2 objective, Nikon Europe B.V.). 3D images were analyzed using the GA3 module of the NIS-element software (Advanced research 6.20.02 version, Nikon). Briefly, the cells and their mitochondrial content were segmented in 3D by thresholding. The ratio of the volume occupied by the mitochondria to the total volume of the cell was then calculated.

### Scanning Electron Microscopy (SEM)

BCG-and Mb3601-infected (2 h at a MOI of 1 for both) cells, as well as non-infected cells, were plated on glass slides and fixed by incubating for 24 h in 1% glutaraldehyde, 4% paraformaldehyde (Sigma) in 0.1 M phosphate buffer (pH 7.2). Samples were washed in PBS and were then fully dehydrated in a graded series of ethanol solutions and dried in hexamethyldisilazane. Finally, samples were coated with 40 Å platinum, with a GATAN PECS 682 apparatus (Pleasanton), before observation under a Zeiss Ultra plus FEG-SEM scanning electron microscope (Oberkochen).

### Transmission Electron Microscopy (TEM)

BCG-and Mb3601-infected (2 h at a MOI of 1 for both) cells, as well as non-infected cells, were fixed by incubating for 24 h in 1% glutaraldehyde, 4% PFA (Sigma) in 0.1 M phosphate buffer (pH 7.2). Samples were then washed in PBS and post-fixed by incubation for 1 h with 2% osmium tetroxide (Agar Scientific). Cells were then fully dehydrated in a graded series of ethanol solutions and propylene oxide. They were impregnated with a mixture of (1∶1) propylene oxide/Epon resin (Sigma) and left overnight in pure resin. Samples were then embedded in Epon resin (Sigma), which was allowed to polymerize for 48 h at 60°C. Ultra-thin sections (90 nm) of these blocks were obtained with a Leica EM UC7 ultramicrotome (Wetzlar). Sections were stained with 2% uranyl acetate (Agar Scientific), 5% lead citrate (Sigma) and observed in a transmission electron microscope (JEOL 1011). For quantitative analysis, ultrastructural features were monitored, with NIS-elements D5 software (Nikon). For the quantification of granules neutrophils, cells of comparable size with a clearly visible nucleus and optimal contrast were selected. Granules were manually counted in five individual cells. Granule size measurements were performed on the same micrographs used for granule quantification. Images were recalibrated prior to measurement by defining the scale bar displayed on each micrograph as the reference distance. Approximately 30 granules per cells were measured in five neutrophils.

## Statistical analysis

Individual data are presented together with the median (Fig. 2b, 2c, 3, 6c, 8g, 8h, Sup. Fig. 1) or mean (Fig. 4b, 4c, 5b) in the figures. Statistical analyses were performed with Prism 10.5.0 software (GraphPad). Analyses were performed on data from two to eight independent experiments. The non-parametric Wilcoxon matched-pairs signed-rank test was used for Fig. 2b, 2c, 3, 4b, 4c, 5b. The non-parametric Mann-Whitney test was used for Fig. 6c. The two-way ANOVA test was used for Fig. 8g and 8h. The *p*-values obtained were interpreted as follows: ns=non-significant; \**p*<0.05; \*\**p*<0.01; \*\*\**p*<0.001, \*\*\*\**p*<0.0001.

**Fig. 1:**
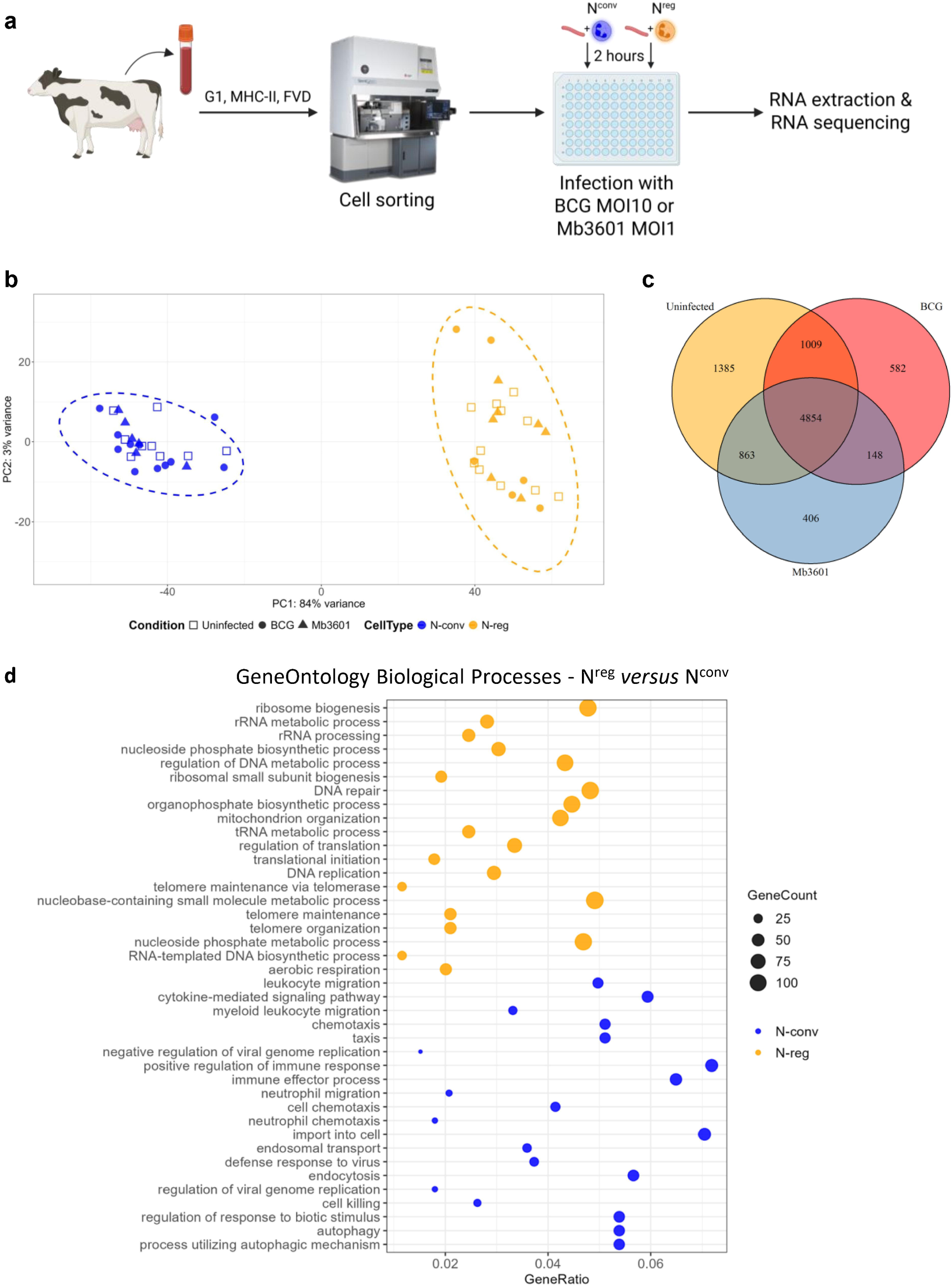
N^reg^ and N^conv^ have different transcriptional profiles. **(a)** Schematic representation of the sequence of steps in the experiment. Cattle blood cells were sorted to separate the two subsets of neutrophils. N^reg^ and N^conv^ were incubated for 2 h either with medium or Mb strains (BCG-WT at a MOI of 10, Mb3601 at a MOI of 1). RNA was then extracted from these samples and sequenced (RNAseq). **(b)** Principal component analysis (PCA) was conducted and the two first dimensions of the PCA plot are shown. **(c)** Venn diagram illustrating the number of genes for each set of conditions, including those common to multiple conditions. **(d)** Dot plot of the top 20 Gene Ontology Biological Process pathways for each neutrophil subtype, for all conditions considered together (4854 differentially expressed genes (DEGs)). Pathways are sorted from top to bottom according to their *p*-values, all of which are significant (*p*<0.05). Orange dots represent the pathways upregulated in N^reg^ and downregulated in N^conv^, whereas blue dots represent pathways that are upregulated in N^conv^ and downregulated in N^reg^. Dot size is proportional to Gene Count. The data shown are for individual samples of neutrophils from *n*=6-11 independent cows, pooled data from 11 independent experiments.

**Fig. 2:**
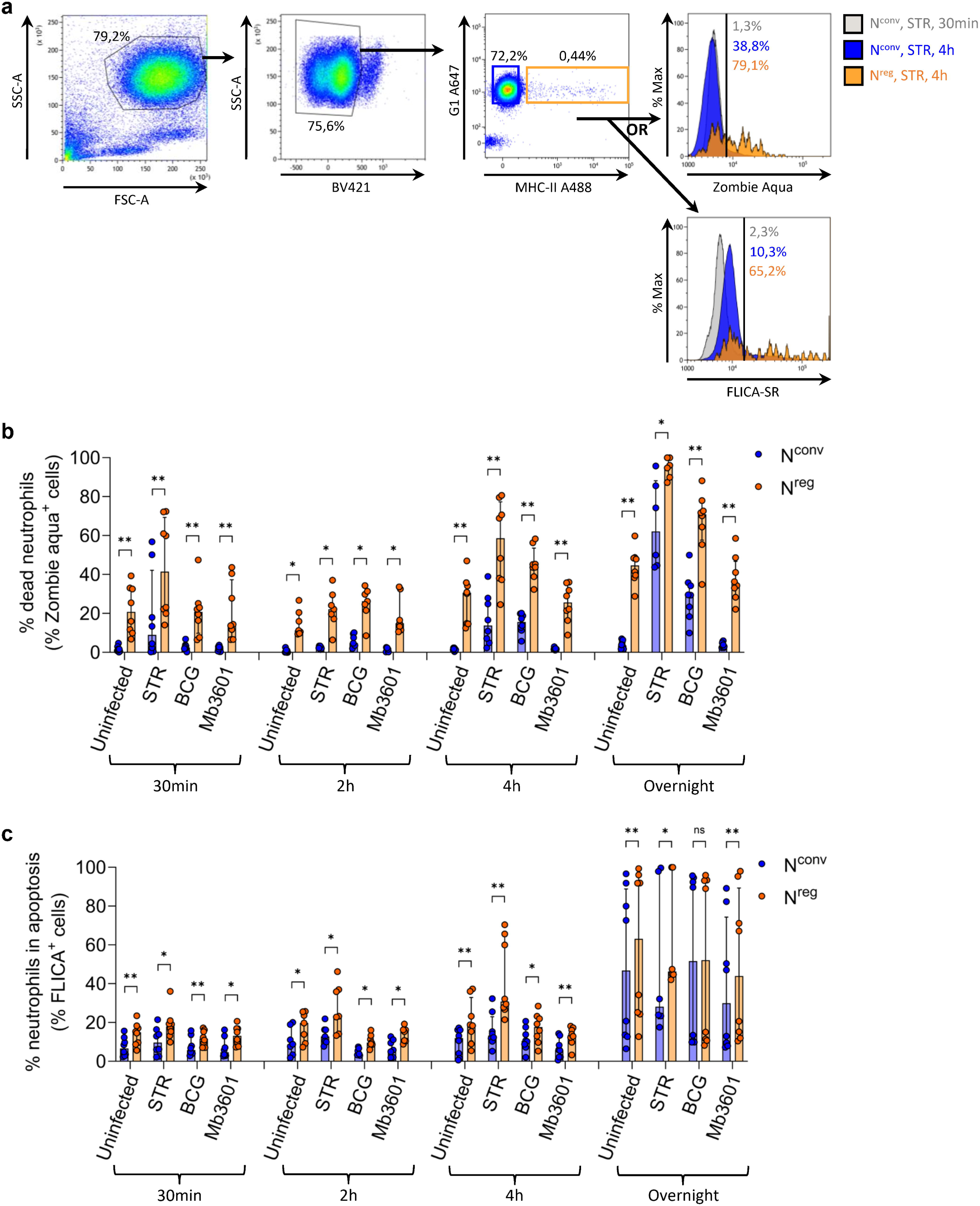
N^reg^ have a higher rate of cell death and die faster than N^conv^. Cattle blood cells were centrifuged and granulocytes were isolated by removing the plasma layer and buffy coat and lysing the red blood cells. Granulocytes were then labeled. **(a)** Gating strategy for quantifying neutrophil death and apoptosis, with data for 1 representative animal shown. Eosinophils were excluded on the basis of their autofluorescence for BV421. Dead blood N^conv^ are defined as SSC-A^high^ FSC-A^high^ BV421^-^ G1^+^ MHC-II^-^ Zombie Aqua^+^ cells, whereas dead blood N^reg^ are defined as SSC-A^high^ FSC-A^high^ BV421^-^G1^+^ MHC-II^+^ Zombie Aqua^+^ cells. Blood N^conv^ undergoing apoptosis are defined as SSC-A^high^ FSC-A^high^ BV421^-^ G1^+^ MHC-II^-^ SR-FLICA^+^ cells, whereas blood N^reg^ undergoing apoptosis are defined as SSC-A^high^ FSC-A^high^ BV421^-^ G1^+^ MHC-II^+^ SR-FLICA^+^ cells. Staurosporine (STR) was used to induce cell death as a positive control. **(b)** After labeling, cells were infected with BCG at a MOI of 10 or Mb3601 at a MOI of 1 or were treated with STR. Cell death rates were assessed by staining with a viability dye (Zombie Aqua). **(c)** After labeling, cells were infected with BCG at a MOI of 10 or Mb3601 at a MOI of 1. Cell apoptosis was assessed with a SR-FLICA polycaspase kit (SR-FLICA) detecting active caspases in neutrophils. **(b-c)** The data shown are the median and interquartile range for neutrophils from *n*=7-8 independent cows, in two independent experiments. ns = non-significant; \**p*<0.05; \*\**p*<0.01 in the non-parametric Wilcoxon matched-pairs signed-rank test.

**Fig. 3:**
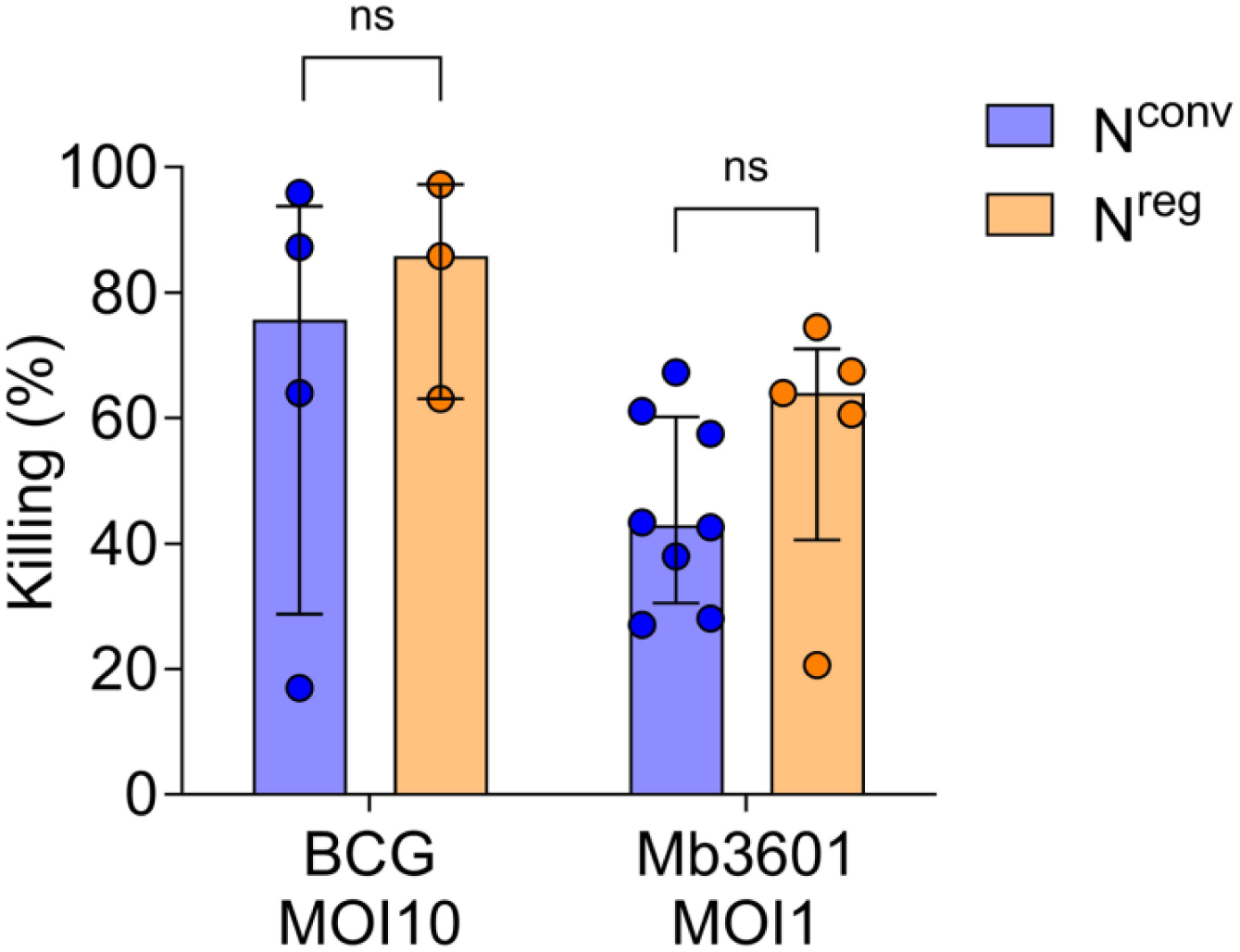
N^reg^ has a capacity to kill BCG and Mb3601 similar to that of N^conv^. After cell sorting, cells were infected with BCG at a MOI of 10 and Mb3601 at a MOI of 1 for 2 h. Both supernatants and lysates were plated and the number of colonies was counted to determine the killing rates as a percentage. The data shown are the median and interquartile range for both subsets of neutrophils from *n*=3-8 independent cows, in 8 independent experiments. ns = non-significant in the non-parametric Wilcoxon matched-pairs signed-rank test.

**Fig. 4:**
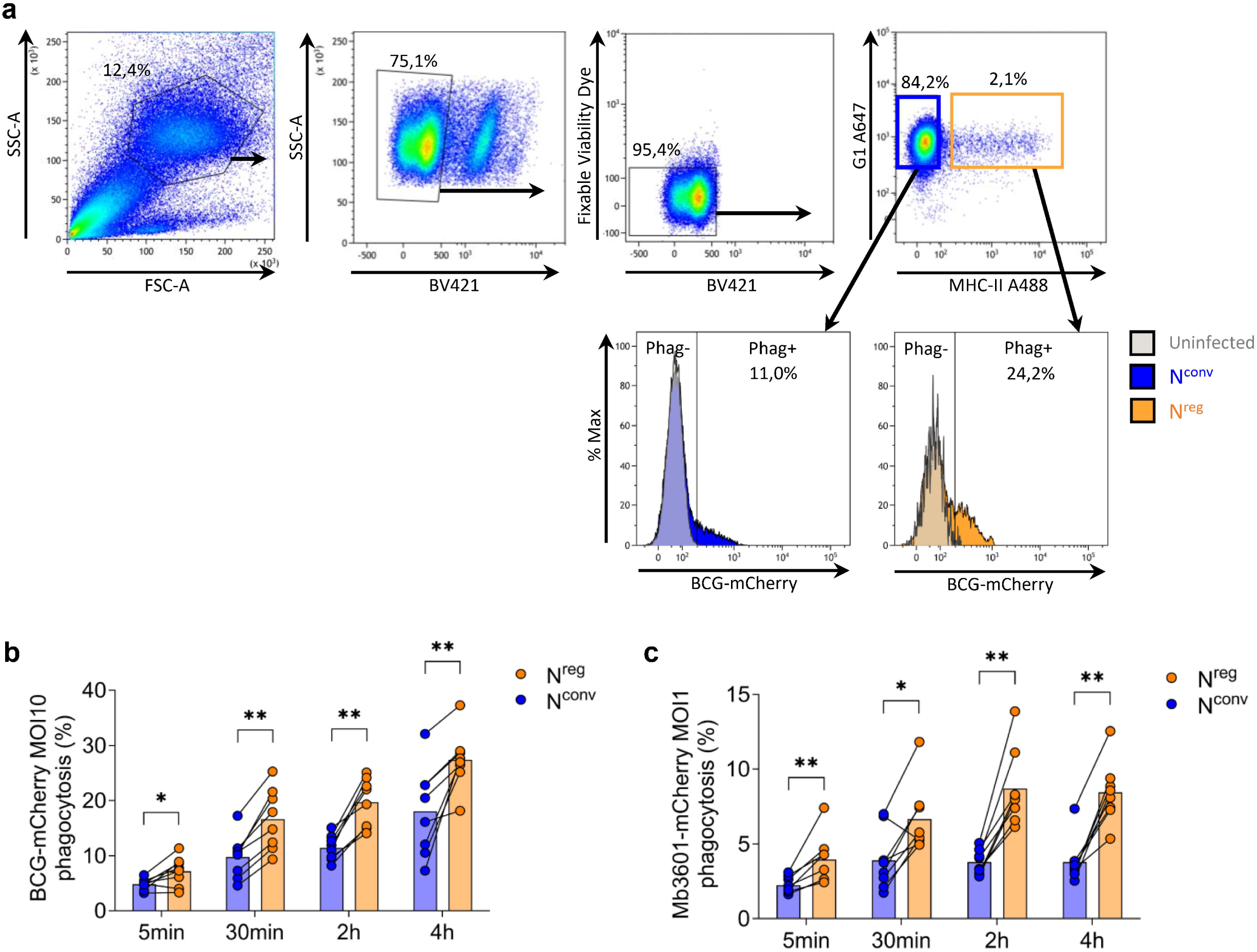
N^reg^ performs phagocytosis more effectively than N^conv^. Granulocytes from cattle blood were treated and labeled as described in Fig. 2. **(a)** Gating strategy for the evaluation of phagocytosis; data for one representative animal shown. Eosinophils were excluded on their autofluorescence for BV421. Viable blood N^conv^ phagocytosing bacteria are defined as SSC-A^high^ FSC-A^high^ BV421^-^ FVD^-^ G1^+^ MHC-II^-^ mCherry^+^ cells, whereas viable blood N^reg^ phagocytosing bacteria are defined as SSC-A^high^ FSC-A^high^ BV421^-^ FVD^-^ G1^+^ MHC-II^+^ mCherry^+^ cells. After labeling, cells were infected with **(b)** BCG-mCherry at a MOI of 10 or **(c)** Mb3601-mCherry at a MOI of 1. The data shown are the mean values and before-after comparisons for the two subsets of neutrophils from *n*=8 independent cows, in 2 independent experiments. \**p*<0.05; \*\**p*<0.01 in non-parametric Wilcoxon matched-pairs signed-rank tests.

**Fig. 5:**
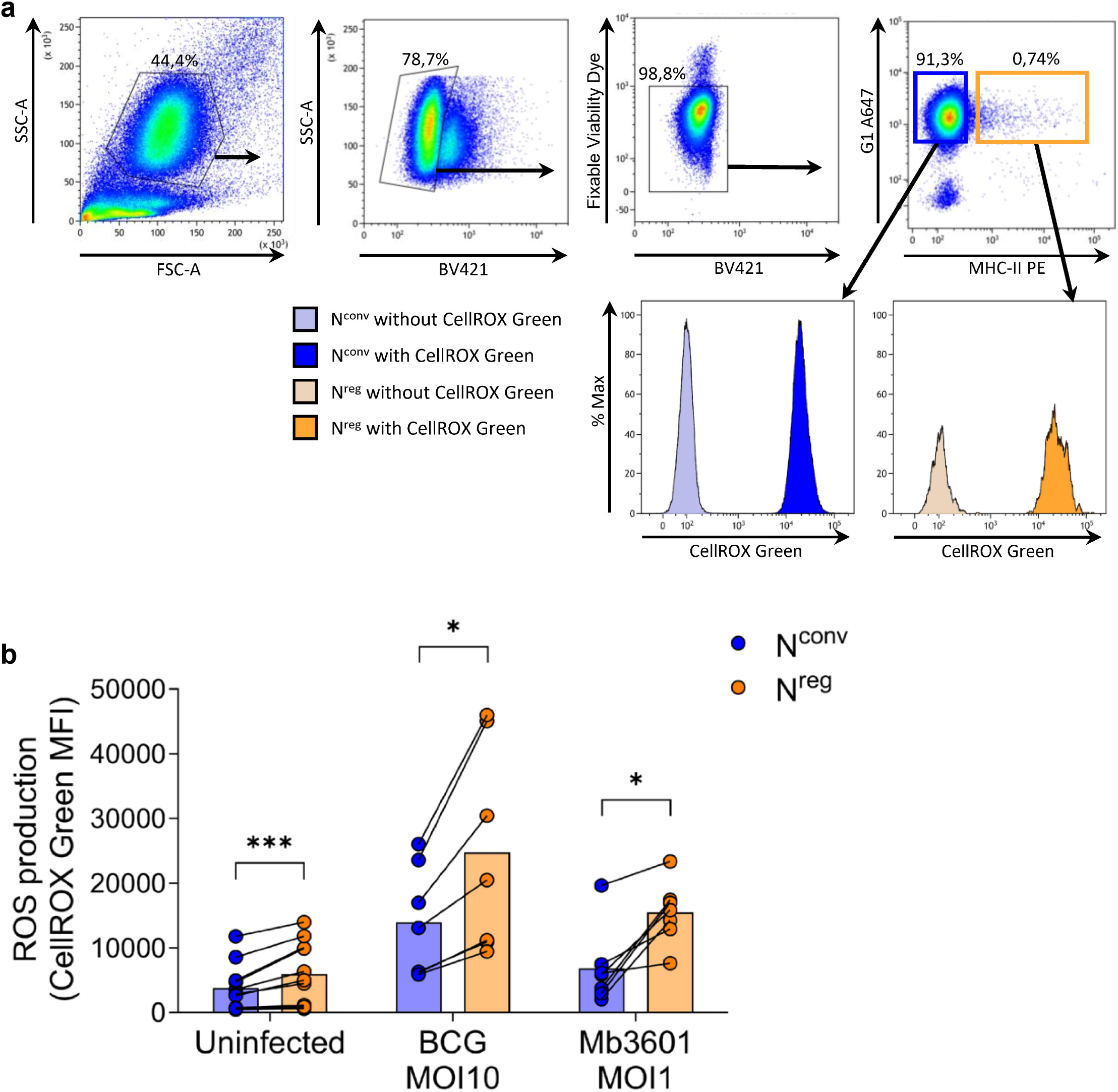
N^reg^ produce higher levels of reactive oxygen species (ROS) than N^conv^. Cattle granulocytes from blood were treated and labeled as described in Fig. 2. **(a)** Gating strategy for the evaluation of ROS production, with data for 1 representative animal shown. Eosinophils were excluded from whole granulocytes on the basis of their autofluorescence for BV421. Viable blood N^conv^ are defined as SSC-A^high^ FSC-A^high^ BV421^-^ FVD^-^ G1^+^ MHC-II^-^ CellROX Green^+^ cells, whereas viable blood N^reg^ are defined as SSC-A^high^ FSC-A^high^ BV421^-^ FVD^-^ G1^+^ MHC-II^+^ CellROX Green^+^ cells. After cell labeling, **(b)** oxidative stress was measured in both neutrophil subpopulations in the presence and absence of infection, over a period of 2 h, with the CellROX Green probe. This probe reacts with certain types of ROS species, such as the hydroxyl radical (HO) and the superoxide anion (O_2_^-^). Mean fluorescence intensity (MFI) was quantified by flow cytometry. The data shown are the mean and before-after comparisons between the two subsets of neutrophils from *n*=7-11 independent cows, in 4 independent experiments. \**p*<0.05; \*\*\**p*<0.001 in non-parametric Wilcoxon matched-pairs signed-rank tests.

**Fig. 6:**
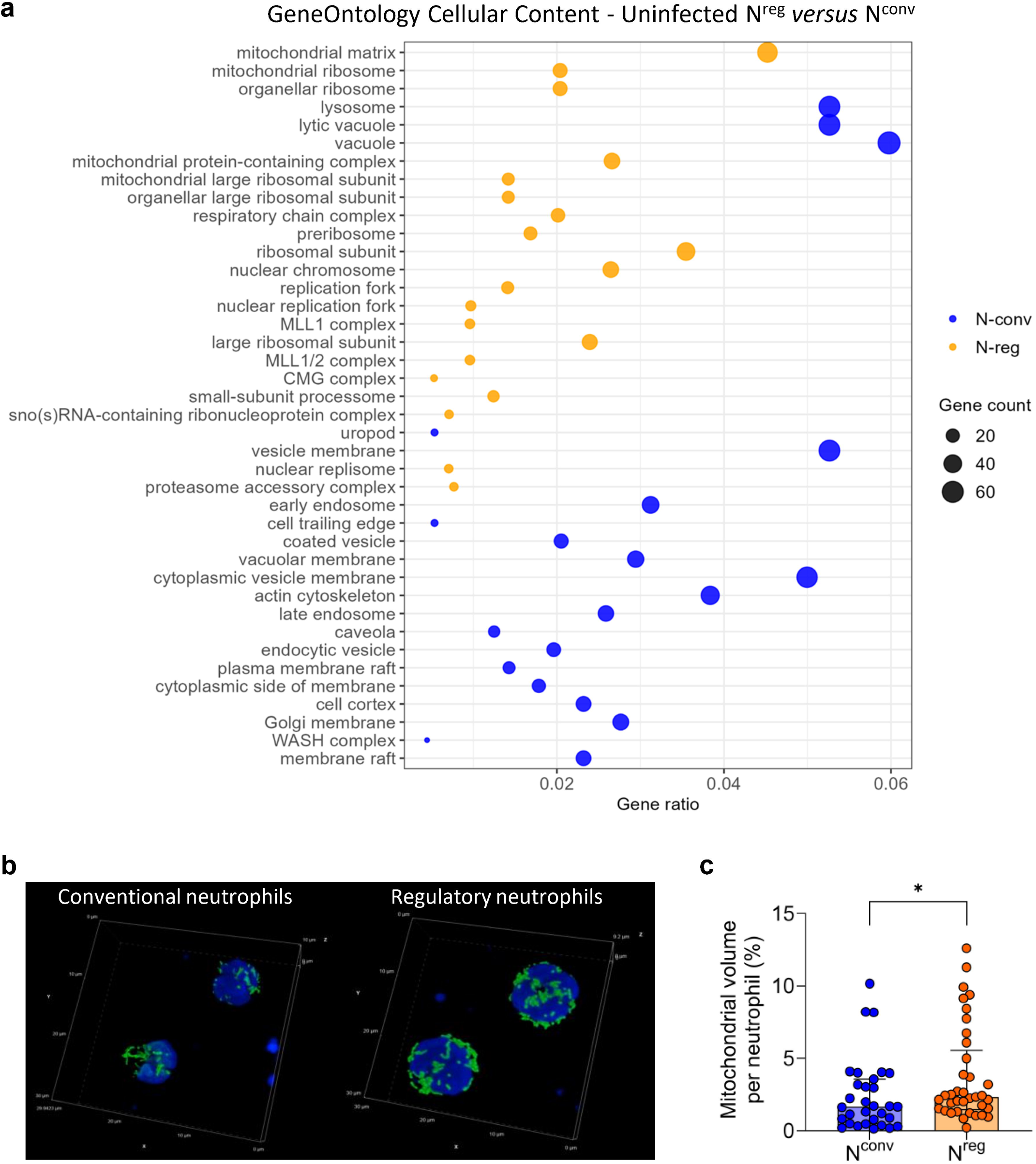
N^reg^ have a higher mitochondrial load than N^conv^. **(a)** Dot plot of the top 20 cellular contents identified for each neutrophil subtype in uninfected conditions (8111 DEG). Pathways are sorted from top to bottom according to their *p*-value, all of which are significant (*p*<0.05). Orange dots represent pathways that are upregulated in N^reg^ and downregulated in N^conv^, whereas blue dots represent pathways that are upregulated in N^conv^ and downregulated in N^reg^. Dot size is proportional to Gene Count. **(b)** Cattle blood N^conv^ and N^reg^ imaged after cell sorting. Nuclei are labeled with Hoechst stain and mitochondria are labeled with Mitotracker Orange CMTMRos, 1 representative animal. **(c)** Mitochondria were quantified with NIS-Element software and GA3 modules. **(a)** The data shown are for individual samples of neutrophils from *n*=6-11 independent cows, pooled data from 11 independent experiments. **(c)** The data shown are for *n*=31-37 neutrophils from 1 cow, 1 experiment. \**p*<0.05 in non-parametric Mann-Whitney test.

## Results

### N^reg^ and N^conv^ have different transcriptional profiles and activate different programs in response to Mb infection

We compared the transcriptomes of N^conv^ and N^reg^ both at steady state and after Mb infection. We purified the two subsets, infected them by incubation for 2 h with avirulent BCG at a MOI of 10, or virulent Mb3601 at a MOI of 1 and conducted a bulk RNA-seq analysis (**Fig. 1a**). An unsupervised principal component analysis (PCA) revealed clear differences between N^reg^ and N^conv^ (**Fig. 1b**) at steady state. The first axis, which accounted for 84% of the variance, separated the two subsets, whereas the second axis, which accounted for only 3% of the variance, did not distinguish clearly between the groups. The strong differential signature between N^conv^ and N^reg^ eclipsed the more subtle effect of bacterial infection in the PCA: uninfected, BCG-infected and Mb3601-infected samples were not clearly separated. We further investigated the impact of Mb infection on the transcription program of the two subsets of neutrophils, by separately comparing the impact of each set of infection conditions on N^reg^ and N^conv^ in the DESeq2 model. We identified 4854 genes commonly modulated across conditions, 582 of which were specifically regulated by BCG infection and 406 by Mb infection (**Fig. 1c**). Using the Gene Ontology knowledgebase, we compiled a list of the top biological pathways in the core set of genes differentially expressed between our two neutrophil subsets (**Fig. 1d**), and then the core sets of genes modulated by infection with the avirulent BCG (**Sup. Fig. 1a**) and virulent Mb3601 strains (**Sup. Fig 1b**). For N^conv^, the biological processes identified were mostly associated with biological processes related to the immune system response and defense (cell migration, cytokines, killing, autophagy, viral immune responses). For N^reg^, the specific biological processes identified were mostly related to metabolic processes and cellular maintenance (cellular biogenesis, protein synthesis, DNA repair and metabolism). Only two pathways were significantly modulated by BCG infection, and both were upregulated in N^reg^: mitotic nuclear division and mitochondrial RNA metabolic processes (**Sup. Fig. 1a**). By contrast, after Mb3601 infection, 20 pathways were upregulated in N^conv^, mostly related to type-I interferon-driven immune responses and T cell activation and proliferation (**Sup. Fig. 1b**). These pathways are representative of the neutrophil-driven dysregulation of the immune response that occurs during active tuberculosis in humans [12]. The other pathways identified were related to stress-induced apoptotic processes, with additional contributions from metabolic and epithelial transport pathways. Mycobacterial infection, thus, clearly induced different transcriptional signatures and activated different biological processes in N^reg^ and N^conv^, particularly after infection with the virulent Mb strain Mb3601.

### Activation results in a shorter lifespan in N^reg^ than in N^conv^

Neutrophils are known to be short-lived, but infection or other danger signals can considerably modify their lifespan [42]. Lifespan may also differ between subsets. We therefore compared the cell death and apoptosis of uninfected and BCG-infected (at a MOI of 10) or Mb3601-infected (at a MOI of 1) N^reg^ and N^conv^ by flow cytometry, with Zombie Aqua as a viability dye and SR-FLICA polycaspase as a marker of apoptosis (**Fig. 2a**). Staurosporine (STR, 2 µM) was used as a positive control for apoptosis and cell death. N^reg^ died significantly more rapidly than N^conv^, 20% of the cells dying during the first 30 minutes of incubation (**Fig. 2b**). Cell death rates increased over time, with approximately 40-70% of N^reg^ dying overnight (> 16 h), versus only 5-35% of N^conv^ in both the presence and absence of infection. As expected, neutrophils were sensitive to STR, which accelerated cell death, beginning after 2 h of incubation. Interestingly, uninfected and infected Mb3601 neutrophils had similar death kinetics, whereas BCG infection induced significantly higher levels of cell death in both subpopulations. Consistent with the observed cell death kinetics, apoptosis rates were higher for N^reg^ than for N^conv^ (**Fig. 2c**). This difference was already visible at 30 min and remained stable for up to 4 h unless STR was added, as expected. Overnight, apoptosis rates approached 50% for both subsets of neutrophils, except for Mb3601-infected, which had lower apoptosis rates. Nevertheless, even for Mb3601 infection, apoptosis rates remained consistently higher for N^reg^ than for N^conv^.

### N^reg^ have a similar level of microbicidal activity but higher levels of phagocytosis and ROS production than N^conv^

We investigated the functions of the two neutrophil subsets in greater detail by first comparing their overall killing activity. We sorted N^reg^ and N^conv^ by flow cytometry and infected them with BCG at a MOI of 10 or Mb3601 at a MOI of 1. We counted CFU both in the supernatant (viable extracellular bacilli) and within neutrophils (viable intracellular bacilli) to determine overall bacterial killing rates. The two subsets of neutrophils had similar abilities to kill non-virulent and virulent strains of Mb (**Fig. 3 and Supp. Fig. 1**). In order to correlate phagocytosis and killing, we infected mixed unsorted neutrophils with recombinant mCherry-BCG strain at a MOI of 10 or mCherry-Mb3601 strain at a MOI of 1 and assessed phagocytosis by flow cytometry at multiple timepoints (**Fig. 4a**). Interestingly, at all timepoints considered, N^reg^ had significantly higher levels of phagocytic activity than N^conv^, regardless of the virulence of the strain used (**Fig. 4b and 4c**). Phagocytosis rates were consistently higher for BCG than for Mb, for both subpopulations of neutrophils. Finally, we assessed ROS production by unsorted N^reg^ and N^conv^ after infection with BCG at a MOI of 10 and Mb3601 at a MOI of 1 for 2 h. We used the CellROX Green probe to measure hydroxyl radical (HO) and superoxide anion (O_2_ ^-^) levels. Mean fluorescence intensity was quantified by flow cytometry (**Fig. 5a**) and directly correlated with ROS production. Strikingly, N^reg^ produced more ROS at baseline than N^conv^. This trend was maintained when neutrophils were infected, with significantly higher levels of ROS production observed for N^reg^ (**Fig. 5b**).

### N^reg^ have higher levels of mitochondrial activity than N^conv^

RNA-seq identified many pathways related to mitochondrial organization and metabolism (**Fig. 1c),** and ROS production was markedly higher in N^reg^ than in N^conv^ (**Fig. 5b**). We therefore decided to investigate the mitochondrion-associated signature in the two subsets. We found that genes involved in mitochondrion-and ribosome-related pathways were strongly enriched in N^reg^, whereas the N^conv^ signature was dominated by vesicles and vacuolar contents (**Fig. 6a**). Using the MitoTracker Orange probe, which binds to active mitochondria, we analyzed and quantified mitochondrial volume by confocal microscopy on sorted neutrophils (**Fig. 6b and 6c**). N^reg^ had a significantly greater mitochondrial volume per cell, suggesting that this subset had larger numbers of mitochondria than N^conv^. These findings are consistent with our observations for ROS production, as mitochondria can generate hydroxyl radicals and superoxide anions, and the levels of these ROS were markedly higher in N^reg^ than in N^conv^.

### N^reg^ and N^conv^ have different membrane and granule ultrastructural organizations

N^reg^ and N^conv^ cannot be distinguished on the basis of their phenotypes on conventional microscopy [25]. We therefore studied these two subtypes in more detail by scanning electron microscopy (SEM) and transmission electron microscopy (TEM). On SEM, N^conv^ were characterized by a predominantly spherical morphology with a smooth, regular cell surface (**Fig. 7a**). The plasma membrane had a uniformly microfolded surface consisting of short, evenly distributed membrane protrusions, resulting in a finely textured appearance. By contrast, N^reg^ had a more heterogeneous morphology, with frequent irregularly shaped or ovoid cells (**Fig. 7b**). The plasma membrane of N^reg^ had a heterogeneous topography, combining deeply folded regions with prominent, thick membrane ruffles. In addition, N^reg^ frequently displayed long, thin membrane extensions, such as filopodium-like structures and tether-or nanotube-like projections, some of which established direct connections between neighboring cells (**Fig. 7b**). Similar ultrastructural differences between N^conv^ and N^reg^ are also observed under infected conditions (**Fig. 7c, 7d, 7e, 7f**). Thus, the two subsets differed markedly in terms of cell shape, membrane organization, and the abundance of extended membrane protrusions, indicating an intrinsic morphological difference between N^conv^ and N^reg^, reflected in differences in surface architectures and membrane remodeling states.

**Fig. 7:**
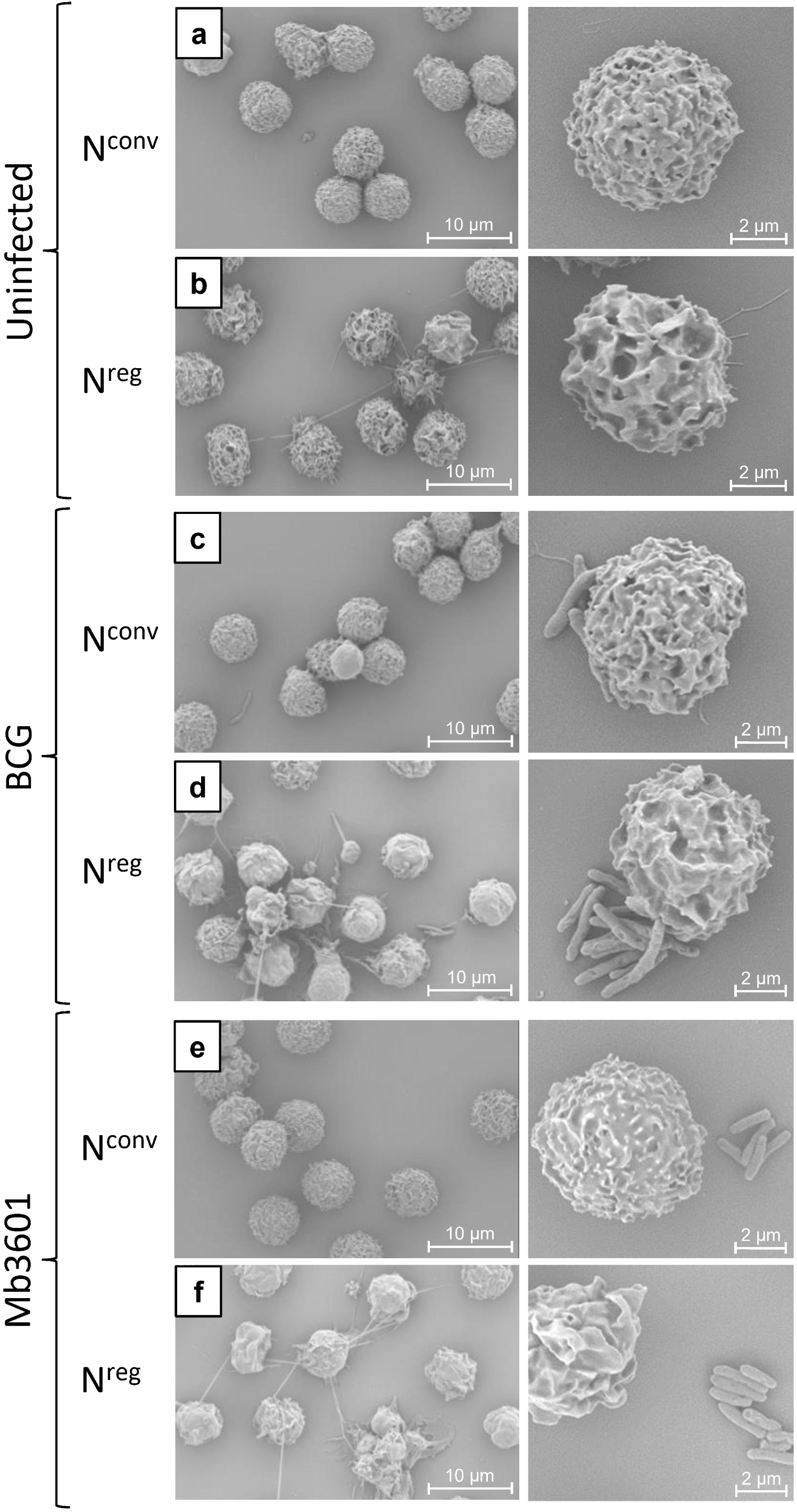
Phenotypic heterogeneity of the two neutrophil subsets on SEM. Sorted cells were infected with BCG or Mb3601 at the MOI of 1 for 2 h. N^reg^ and N^conv^ were fixed and prepared for SEM acquisition. Images of uninfected **(a)** N^conv^ and **(b)** N^reg^, BCG-infected **(c)** N^conv^ and **(e)** N^reg^, and Mb3601-infected **(d)** N^conv^ and **(f)** N^reg^. The data shown are for 5 neutrophils from the same cow, from 1 representative experiment out of 3 experiments.

**Fig. 8:**
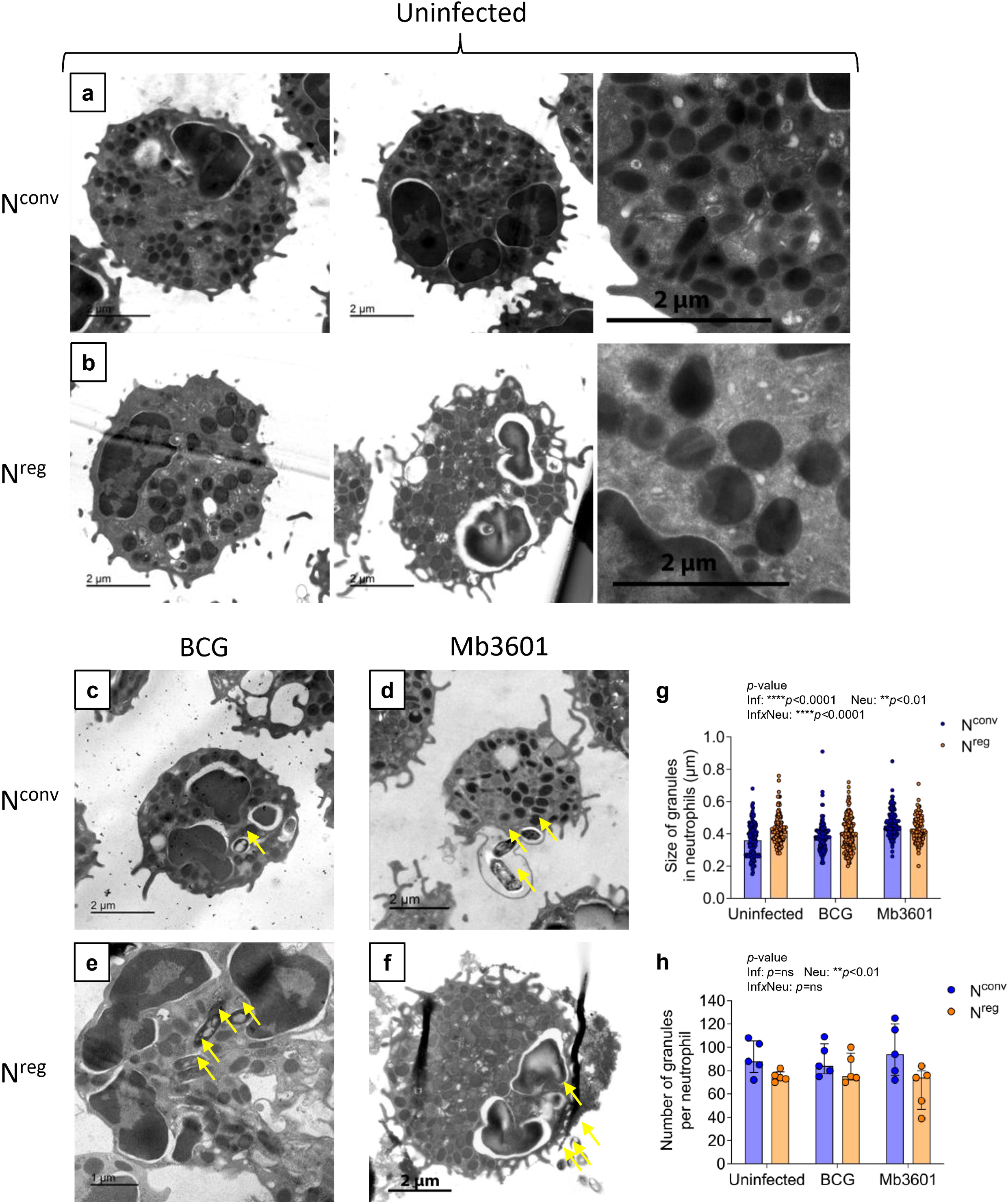
Phenotypic heterogeneity of the two neutrophil subsets in TEM. The sorted cells were infected with BCG or Mb3601 at the MOI of 1 for 2 h. N^reg^ and N^conv^ were fixed and prepared for TEM acquisition. Images of uninfected **(a)** N^conv^ and **(b)** N^reg^, BCG-infected **(c)** N^conv^ and **(e)** N^reg^, and Mb3601-infected **(d)** N^conv^ and **(f)** N^reg^. Mycobacteria are indicated by yellow arrows. Quantitative analysis of **(g)** granule size and **(h)** granule number for each neutrophil subset in the presence and absence of infection. The data shown are the medians for 5 neutrophils from the same cow, pooled from *n*=2 independent cows in 2 independent experiments out of 3 experiments. ns = non-significant; \*\*\**p*<0.001; \*\*\*\**p*<0.0001 in two-way ANOVA test.

In TEM analysis, N^conv^ had a finely textured and densely packed cytoplasm **(Fig. 8a)**. The plasma membrane of N^conv^ had few ultrastructural irregularities, with only short protrusions and modest membrane ruffling. The cortical cytoplasm appeared relatively uniform, without large membrane invaginations or extended cytoplasmic projections, consistent with the SEM observations. The plasma membrane of N^reg^ frequently had pronounced ultrastructural irregularities, including membrane ruffles, protrusions, and occasional thin cytoplasmic extensions **(Fig. 8b)**. The peripheral cytoplasm appeared less uniform, with localized thinning and regions suggestive of enhanced membrane remodeling or granules exocytosis, consistent with the SEM findings. In both the N^conv^ and N^reg^ subpopulations, and following exposure to BCG (**Fig. 8c and 8e**) or Mb3601 (**Fig. 8d and 8f**), direct neutrophil-bacterium contacts, phagocytic events, and intracellular bacteria enclosed within membrane-bound vacuoles were observed. Some N^reg^ had ultrastructural features suggestive of an increase in basal activity, including pronounced membrane remodeling and cytoplasmic or plasma membrane areas consistent with early or partial degranulation.

N^conv^ displayed an evenly distributed granule population with a markedly heterogeneous morphology: a mixture of round, ovoid, and elongated (elliptical) shapes **(Fig. 8a)**. By contrast, N^reg^ were characterized by the presence of less numerous and larger cytoplasmic granules, which appeared more homogeneous in shape, most being regular and round **(Fig. 8b)**. However, the granules were unevenly distributed within the cytoplasm, due in part to the presence of large vacuolar or vesicular compartments resulting in areas of relative granule depletion. Quantitative TEM analysis revealed distinct differences in granule size **(Fig. 8g)** and abundance **(Fig. 8h)** between N^conv^ and N^reg^, depending on infection status. In the absence of infection, N^reg^ contained significantly larger granules, but fewer granules per cell compared to N^conv^. Following infection, N^conv^ retained a higher granule count and exhibited a significant increase in mean granule size, particularly in response to the virulent Mb3601 strain. In contrast, N^reg^ showed a reduction in granule number, while granule size remained relatively large. This divergence suggests differential remodeling of the granular compartment in response to the virulent Mb3601 strain.

## Discussion

In humans and mouse models, neutrophils are considered to be key cells in the pathophysiology of tuberculosis, exerting either protective or deleterious effects depending on the stage and severity of the disease [28,43]. Over the last decade, neutrophil heterogeneity has been widely described in homeostatic conditions, cancers, inflammation and infectious diseases. It is now well established that different neutrophil subsets promote pro-and anti-inflammatory responses and modulate both innate and adaptive immunity.

Little is known about the role of bovine neutrophils during Mb infection. As in mice, sequential studies have revealed a prominent neutrophil component of the host response to Mb in early-stage lesions [44,45]. Wang *et al*. detected a neutrophil gene signature in Mb-infected cattle [46] but *in vitro*, depending on the cytokine environment, neutrophils did not appear to be able to eliminate the bacteria [47]. Another study suggested a deleterious role of neutrophils through strong IL-17 production, contributing to the pro-inflammatory response in lung lesions, and the production of arginase, decreasing Mb killing [48].

Following the discovery of the two neutrophil populations, N^conv^ and N^reg^, circulating in cattle blood [25], this study demonstrates that these two populations have very different features, both at steady state and following infection with Mb *in vitro*, highlighting the need to reconsider the roles of these two bovine neutrophil subsets in the pathophysiology of bTB.

We found that N^conv^ and N^reg^ had different transcriptomic signatures. In N^conv^, an enrichment was observed in the expression of genes involved in the immune functions expected for classical neutrophils, whereas the N^reg^ transcriptomic signature was mostly related to metabolic functions and cellular maintenance. Strikingly, the two neutrophil subsets also had different responses to the virulent Mb3601 strain, with only N^conv^ displaying an upregulation of pathways classically associated with disease activation and poor outcomes in TB, such as the type I interferon pathway [12,49–51]. Maerzdorf *et al.* described functional correlations between pathogenesis-driven gene expression signatures in human TB, such as immune inflammatory responses (involving the JAK-STAT pathway), apoptosis, responses to bacterial molecular patterns and the NF-kappa B cascade; all these key pathways are upregulated in N^conv^ and downregulated in N^reg^, suggesting that N^conv^ is probably the subset responsible for the gene signature classically described in human TB [52]. In an original *ex vivo* bovine lung explant model, we recently observed the local induction by Mb infection of a strong type I interferon response and a neutrophil recruitment signature [53]. Interestingly, Mb exploits the type I IFN signaling pathway for its own survival in bovine macrophages [54,55]. It is unknown whether neutrophils contribute to this IFN type I signature during Mb infection in cattle, as demonstrated in human patients with active TB patients [12].

The ability of neutrophils to kill Mtb or Mb remains a matter of debate [17]. Some authors have reported an impairment of killing by neutrophils in mice and humans [56,57] whereas others have reported effective killing by neutrophils [58,59]. Wang and coworkers investigated the effect of Mb infection on bovine neutrophils obtained from the blood by gradient centrifugation [47]. They showed that these neutrophils were able to kill *M. smegmatis* but not a virulent Beijing strain of Mb. Our data show clear bactericidal activity for both N^conv^ and N^reg^ against both *M. bovis* BCG and the French virulent strain Mb3601. This discrepancy may be due to the sorting of neutrophils in our study, a process ensuring a high purity of both subsets, facilitating their study but also activating them. We cannot exclude the possibility that the remaining Mb bacilli present within the neutrophils were able to replicate, increasing CFU counts at later time points.

N^reg^ share functional similarity to granulocytic myeloid-derived suppressor cells (G-MDSCs), as they are able to suppress T cell proliferation under steady state conditions [25]. G-MDSCs have been described as a permissive niche for Mtb replication, thereby facilitating immune evasion by the pathogen [60]. MDSCs have also been shown to support Mtb replication in the *in vitro* granuloma model, and to alter the structure of the granuloma [61]. They are also associated with disease progression and greater lethality in TB [62]. By contrast, N^reg^ display no such behavior, as they have been shown to kill BCG and Mb3601 *in vitro*.

Neutrophils are widely considered to be professional phagocytes, able to engulf various pathogens, foreign particles or even cell debris. In 2021 and 2023, Rambault *et al* [25,27] reported that both neutrophil subsets could engulf *E. coli* bioparticles and whole *E. coli*-GFP by phagocytosis, with N^reg^ making better phagocytes than N^conv^. We also found that both neutrophil subsets were also able to take up Mb by phagocytosis; however, N^reg^ again had a significantly greater phagocytic capacity than N^conv^, at 30 min and after up to 4 h of infection. The rapid and enhanced phagocytic activity of N^reg^ is not, therefore, dependent on the bacterial strain, with this activity instead being a general, non-specialized property of this subtype. Remarkably, RNAseq analysis based on Gene Ontology Biological Process or Cellular Component identified various pathways and organelles associated with phagocytosis in N^conv^ only, with certain terms: autophagy, endocytosis, lysosome, early and late endosome, and vacuole. These data suggest that, upon initial contact with Mb, N^reg^ are already primed for phagocytosis, accounting for their greater phagocytic capacity at early time points. Interestingly, we previously reported higher levels of expression of CD11b [25], an important target for mycobacterial entry, on N^reg^ [63].

Reactive oxygen species (ROS) are known to be able to kill pathogens, but their role in eliminating mycobacteria remains a matter of debate. Some *in vitro* studies have suggested that ROS production is required for the killing of Mtb [64–66], whereas others have reported resistance to oxidative stress in Mtb in mice[67] or impaired neutrophil killing by ROS in humans [57]. ROS are also known to act as potent immunomodulators in multiple contexts, influencing adaptive immunity through T and B cells [68–71]. N^reg^ and N^conv^ displayed similar levels of bacterial killing but N^reg^ displayed a significantly larger increase in ROS production on infection than N^conv^. As N^reg^ have a unique ability to regulate T cells [25,26] and ROS are known to be major compounds produced by granulocytic MDSCs to alter the T cell receptor [72], we hypothesize that the higher levels of ROS production in N^reg^ not only help to kill Mb, but that they also have immunomodulatory effects on other immune cells, such as T cells, especially during the later stages of TB.

N^reg^ display a significant increase in ROS production accompanied by the activation of numerous pathways related to metabolism and mitochondrial organization. As mitochondria are also a major source of intracellular ROS, we investigated the mitochondrial content of this population to compare N^reg^ and N^conv^ further. The mitochondrial compositions of neutrophil subsets remain poorly characterized. Rice *et al*., 2018, identified two subsets of neutrophils in mice with and without cancers: Ly-6G^+^/c-Kit^+^/CXCR2^-^ cells, which are less mature and more oxidative than the other subset, Ly-6G^+^/c-Kit^-^/CXCR2^+^ neutrophils [73]. Interestingly, c-Kit^+^/CXCR2^-^ neutrophils had a mitochondrion-rich phenotype, with enhanced free radical production, especially under glucose-limited conditions. This subtype has several features in common with our N^reg^, which have a higher mitochondrial load, as demonstrated by confocal microscopy, and higher levels of ROS production than N^conv^. Furthermore, RNAseq analysis revealed an upregulation of the *kit* gene (+3.075 log_2_FC) and a downregulation of the *cxcr2* gene (-1.58 log_2_FC) in N^reg^, consistent with these findings.

However, even though c-Kit^+^/CXCR2^-^ neutrophils are considered to be immature, the N^reg^ we studied here had a highly segmented nuclear morphology similar to that of mature neutrophils [25].

In addition to these observations, the high levels of ROS production observed in N^reg^ may account for the higher rates of cell death detected in this subset. Indeed, enhanced ROS levels are associated with cell damage and impaired viability, leading to the activation of cell death-related pathways [74,75]. This study provides evidence of different death dynamics between neutrophil subsets, particularly as concerns apoptosis. Our data indicate that N^reg^ not only die more rapidly than N^conv^, but that they undergo apoptosis at a higher rate in both steady-state and infection conditions, suggesting an intrinsic predisposition towards lower survival. In our RNAseq analysis comparing the two subsets, only N^reg^ displayed an enrichment in a few significant pathways related to telomeres and telomerase that might explain higher death rates more generally. However, when neutrophils were infected with Mb3601, N^conv^ displayed an enrichment in pathways related to the up-or downregulation of apoptosis (**Fig. S1b**). No such effect was observed after infection with BCG. This may be due to the virulence of Mb3601, resulting in a greater capacity of this strain to modulate host responses. Neutrophil death occurred in a pathogen-dependent manner: BCG infection accelerated cell death, whereas Mb3601 infection appeared to delay it, highlighting the capacity of mycobacterial strains to influence cell lifespan differentially [76]. Apoptosis is one of the principal pathways contributing to neutrophil death, but it is unlikely to be sufficient to account entirely for the observed differences between neutrophil subsets. Other types of cell death, such as necrosis or NETosis, may also be involved and could have important functional consequences for inflammation and host-pathogen interactions, especially in the context of TB [77,78]. Indeed, NETosis is also a crucial process in TB, with high NET levels associated with a poor disease outcome [79,80]. We attempted to investigate NETosis in both N^reg^ and N^conv^ by confocal microscopy, after labeling NETs with antibodies directed against myeloperoxidase, neutrophil elastase or citrullinated histone H3. We observed rare NETs *in vitro* (data not shown), probably not entirely representative of what has been seen *in vitro* in other models [81] or *in vivo* [82]. Cell sorting may account for the low frequency of NET formation *in vitro*, as this procedure can activate neutrophils, altering their capacity to undergo NETosis. This does not necessarily exclude the possibility of such events occurring *in vivo*. However, our RNAseq analysis with the KEGG database revealed an upregulation of the neutrophil extracellular trap pathway in N^conv^ (**Sup. Material 1**), suggesting that disease exacerbation *in vivo* may be dependent only on NETs from N^conv^. Further investigations of these alternative death pathways are now required to improve our understanding of the fate of neutrophils and the regulatory role of these cells in bTB.

We were unable to identify any morphological differences between N^reg^ and N^conv^ on conventional microscopy [25]. However, SEM and TEM revealed remarkable differences in morphology, relating to cell shape, membrane and granule organization, between N^conv^ and N^reg^. These ultrastructural differences may underlie different basal functional states in neutrophils. N^conv^ have features consistent with a structurally stable quiescent state, whereas N^reg^ might have a lower activation threshold and greater membrane plasticity, potentially conferring a greater propensity for rapid cell-cell and cell-bacteria interactions. We noted important structural differences between the two subsets, including the presence of thin, elongated filopodium-or nanotube-like structures. Such structures have been described in macrophages [83,84], but have rarely been described in neutrophils. We cannot confirm their exact nature, but we hypothesize that these protrusions may correspond to tunneling nanotubes or proteoglycofili [85], involved in communication between cells or pathogen neutralization. Differences in granule size and number between the two neutrophil subsets further support their phenotypic divergence. The neutrophil granules of cattle differ from those described in mice and humans, as cattle have a specific type of granule known as azurophilic granules, and a third, as yet unnamed, type of granule [86]. It was not possible to distinguish these types of granules by electron microscopy in our neutrophil subsets, but transcriptomic analysis provided further insight. Reactome pathway analysis (**Sup. Material 1**) revealed a significant enrichment in the neutrophil degranulation pathway in N^conv^, suggesting a greater antimicrobial effect of degranulation in this subset. This hypothesis was supported by the upregulation of several granule-associated genes in N^conv^, including *elane*, *ltf*, *s100a9*, *mmp25* and *olfm4*. However, N^reg^ also expressed genes related to granules: *bpi*, *cybb*, *lyz*, *defb10*, *defb4*, and members of the serpin family (*serpinb1*, *serpinb2*, and *serpinb6*). Together, these findings point to the existence of two different granule-related programs in these two subsets of neutrophils. Further investigations, including granule-focused proteomics analyses in particular, will be required to determine how these differences translate into functional outcomes and whether they are modulated by Mb infection. Neutrophils have been shown to act as a source of antimicrobial agents potentiating the killing of Mtb by macrophages through the delivery of their granule contents [87], but the survival of Mtb within macrophages has been shown to be enhanced following the ingestion of neutrophils containing this bacterium [88]. As N^conv^ and N^reg^ have different granule contents, this crosstalk between neutrophils and macrophages should be investigated further, taking the heterogeneity of neutrophils into account.

In conclusion, our study demonstrates that bovine N^reg^ and N^conv^ are very different types of neutrophils programmed to perform different functions. Following infection with Mb, these two subsets had clearly different transcriptional profiles, but similar bactericidal capacities. N^reg^ displayed accelerated cell death, enhanced phagocytic activity, an increase in ROS production, and a higher mitochondrial load than N^conv^. Electron microscopy-based ultrastructural analyses revealed phenotypic divergences between the two subsets, particularly in terms of cell morphology, membrane organization and granular content. Together, these findings provide the first functional characterization of bovine neutrophil subsets during Mb infection and highlight a new layer of complexity in the functional diversity of neutrophil subsets in TB. Nevertheless, these neutrophil-Mb interactions relate exclusively to the early stage of the disease, and further *in vitro* studies on later stage of bTB (granuloma) or *in vivo* studies will be required to gain a full understanding of the role of N^reg^ in bTB, including their contribution to host immune responses, disease progression and pathological effects on tissues. Elucidation of the specific role of bovine neutrophil subsets in bTB pathophysiology may pave the way for the discovery of new biomarkers, which are urgently needed to improve the management of this costly disease.

## Statements

## Supporting information

SupMaterial1

SupFig1

SupFig2

## Acknowledgement

We warmly thank the UE-PAO experimental unit for blood sampling and animal care. We thank Alix Sausset for her assistance with neutrophil sorting. The Mb3601 strain was obtained from Dr. Maria-Laura Boschiroli (ANSES, French bTB reference laboratory). We thank the Microscopy Facility (US61 ASB) of Tours University and CHRU of Tours (http://microscopies.med.univ-tours.fr) for technical support. We also thank Axelle Guillet and Pauline Courroussé, who performed their Master 1 and professional Bachelor’s degree projects, respectively, in our team and helped with some of the experiments.

## Statement of ethics

Blood sampling was approved by the Ethics Committee for Animal Experimentation Val-de-Loire (CEEA VdL), registered with the French National Committee for Animal Experimentation (APAFIS no. 202007241825481.V2), and complied with EU regulation 2010/63.

## Author Contributions

M Saint-Vanne: conceptualization, formal analysis, investigation, validation, visualization, methodology, and writing — original draft, review, and editing.

B Bounab: conceptualization, software, data curation, formal analysis, validation, investigation, visualization, methodology, and writing — review and editing.

S Eymieux: methodology, resources, formal analysis, validation, visualization and writing — review and editing.

E Perdriau: methodology, investigation, formal analysis, validation and visualization.

F Carreras: methodology and investigation R Roullier: investigation

Y Le Vern: methodology and investigation

J Pichon: methodology, formal analysis and visualization. E Doz-Deblauwe: writing—review and editing.

P Germon: formal analysis and writing — review and editing.

N Winter: conceptualization, and writing — original draft, review, and editing.

A Remot: conceptualization, formal analysis, investigation, supervision, funding acquisition, visualization, validation, methodology, project administration, and writing — original draft, review, and editing.

## Data Availability Statement

**Sup. Fig. 1: Transcriptomic profiles of N^reg^ and N^conv^ following infection with BCG or Mb3601.** Cattle blood cells were sorted to separate the two subsets of neutrophils. N^reg^ and N^conv^ were incubated for 2 h with either medium or one of the Mb strains (BCG-WT at a MOI of 10 or Mb3601 at a MOI of 1). RNA was then extracted from the samples and used for RNAseq analysis. **(a)** Dot plot of the top Gene Ontology Biological Process pathways identified for BCG-infected neutrophils (582+148 DEGs). **(b)** Dot plot of the top Gene Ontology Biological Process pathways for Mb3601-infected neutrophils (406+148 DEGs). Pathways are sorted from top to bottom according to their *p*-values, all of which are significant (*p*<0.05). Orange dots represent pathways that are upregulated in N^reg^ and downregulated in N^conv^, whereas blue dots represent pathways that are upregulated in N^conv^ and downregulated in N^reg^. Dot size is proportional to Gene Count. The data shown correspond to individual samples of neutrophils from *n*=6-11 independent cows, in 11 independent experiments.

**Sup. Fig. 2: CFUs retrieved from lysate and supernatant in the killing assay.** The sorted cells were infected with BCG (MOI 10) or Mb3601 (MOI 1) for 2 h. Both supernatants and lysates were plated and counted to determine the percent killing. The data shown are the medians and interquartile ranges of the subsets of neutrophils from *n*=3-8 independent cows, in 8 independent experiments.

**Sup. Material 1: Comparative pathway analysis of neutrophil subtypes across multiple conditions.** Table of 20 top enriched pathways in neutrophil subtypes using Gene Ontology (Biological Process (BP) and Cellular Component (CC)), Reactome or KEGG databases for each neutrophil subtype across all condition combined (sheet 1) or sorted by condition (sheet 2). Pathways are sorted from top to bottom according to their p-value, all of which are significant (p<0.05). Data represent individual samples of neutrophils from n=6-11 independent cows, pooled data from 11 independent experiments.

## References

1. Conlan, A. J. K., Vordermeier, M., De Jong, M. C. & Wood, J. L. The intractable challenge of evaluating cattle vaccination as a control for bovine Tuberculosis. eLife 7, e27694 (2018).

2. Hénaux, V. et al. Sensitivity of bovine tuberculosis surveillance through intradermal tests in cattle in France: An evaluation of different scenarios. Prev. Vet. Med. 191, 105364 (2021).

3. Michelet, L., De Cruz, K., Tambosco, J., Hénault, S. & Boschiroli, M. L. Mycobacterium microti Interferes with Bovine Tuberculosis Surveillance. Microorganisms 8, 1850 (2020).

4. Tuberculose bovine_: la situation en France. Ministère de l’Agriculture, de l’Agro-alimentaire et de la Souveraineté alimentaire https://agriculture.gouv.fr/tuberculose-bovine-la-situation-en-france.

5. Bouchez-Zacria, M., Courcoul, A. & Durand, B. The Distribution of Bovine Tuberculosis in Cattle Farms Is Linked to Cattle Trade and Badger-Mediated Contact Networks in South-Western France, 2007–2015. Front. Vet. Sci. 5, 173 (2018).

6. Richomme, C. et al. Exposure of Wild Boar to Mycobacterium tuberculosis Complex in France since 2000 Is Consistent with the Distribution of Bovine Tuberculosis Outbreaks in Cattle. PLoS ONE 8, e77842 (2013).

7. Alfonseca-Silva, E., Hernández-Pando, R. & Gutiérrez-Pabello, J. A. Mycobacterium bovis-infected macrophages from resistant and susceptible cattle exhibited a differential pro-inflammatory gene expression profile depending on strain virulence. Vet. Immunol. Immunopathol. 176, 34–43 (2016).

8. Magee, D. A. et al. Innate cytokine profiling of bovine alveolar macrophages reveals commonalities and divergence in the response to Mycobacterium bovis and Mycobacterium tuberculosis infection. Tuberc. Edinb. Scotl. 94, 441–450 (2014).

9. Jensen, K. et al. Variation in the Early Host-Pathogen Interaction of Bovine Macrophages with Divergent Mycobacterium bovis Strains in the United Kingdom. Infect. Immun. 86, e00385–17 (2018).

10. Martineau, A. R. et al. Neutrophil-mediated innate immune resistance to mycobacteria. J. Clin. Invest. 117, 1988–1994 (2007).

11. Eum, S.-Y. et al. Neutrophils Are the Predominant Infected Phagocytic Cells in the Airways of Patients With Active Pulmonary TB. Chest 137, 122–128 (2010).

12. Berry, M. P. R. et al. An interferon-inducible neutrophil-driven blood transcriptional signature in human tuberculosis. Nature 466, 973–977 (2010).

13. Pesenti, L. et al. Neutrophils Display Novel Partners of Cytosolic Proliferating Cell Nuclear Antigen Involved in Interferon Response in COVID-19 Patients. J. Innate Immun. 17, 154–175 (2025).

14. Lombard, R. et al. IL-17RA in Non-Hematopoietic Cells Controls CXCL-1 and 5 Critical to Recruit Neutrophils to the Lung of Mycobacteria-Infected Mice during the Adaptive Immune Response. PloS One 11, e0149455 (2016).

15. Seiler, P. et al. Early granuloma formation after aerosol Mycobacterium tuberculosis infection is regulated by neutrophils via CXCR3-signaling chemokines. Eur. J. Immunol. 33, 2676–2686 (2003).

16. Yang, C.-T. et al. Neutrophils exert protection in the early tuberculous granuloma by oxidative killing of mycobacteria phagocytosed from infected macrophages. Cell Host Microbe 12, 301–312 (2012).

17. Lowe, D. M., Redford, P. S., Wilkinson, R. J., O’Garra, A. & Martineau, A. R. Neutrophils in tuberculosis: friend or foe? Trends Immunol. 33, 14–25 (2012).

18. Muefong, C. N. & Sutherland, J. S. Neutrophils in Tuberculosis-Associated Inflammation and Lung Pathology. Front. Immunol. 11, 962 (2020).

19. Garley, M. & Jabłońska, E. Heterogeneity Among Neutrophils. Arch. Immunol. Ther. Exp. (Warsz.) 66, 21–30 (2018).

20. Ng, L. G., Ostuni, R. & Hidalgo, A. Heterogeneity of neutrophils. Nat. Rev. Immunol. 19, 255–265 (2019).

21. Nicolás-Ávila, J. Á., Adrover, J. M. & Hidalgo, A. Neutrophils in Homeostasis, Immunity, and Cancer. Immunity 46, 15–28 (2017).

22. Loh, W. & Vermeren, S. Anti-Inflammatory Neutrophil Functions in the Resolution of Inflammation and Tissue Repair. Cells 11, 4076 (2022).

23. Jones, H. R., Robb, C. T., Perretti, M. & Rossi, A. G. The role of neutrophils in inflammation resolution. Semin. Immunol. 28, 137–145 (2016).

24. Bassel, L. L. & Caswell, J. L. Bovine neutrophils in health and disease. Cell Tissue Res. 371, 617–637 (2018).

25. Rambault, M. et al. Neutrophils Encompass a Regulatory Subset Suppressing T Cells in Apparently Healthy Cattle and Mice. Front. Immunol. 12, 625244 (2021).

26. Doz-Deblauwe, E. et al. Dual neutrophil subsets exacerbate or suppress inflammation in tuberculosis via IL-1β or PD-L1. Life Sci. Alliance 7, e202402623 (2024).

27. Rambault, M. et al. Neutrophils expressing major histocompatibility complex class II molecules circulate in blood and milk during mastitis and show high microbicidal activity. J. Dairy Sci. 106, 4245–4256 (2023).

28. Lyadova, I. V. Neutrophils in Tuberculosis: Heterogeneity Shapes the Way? Mediators Inflamm. 2017, 1–11 (2017).

29. Abadie, V. et al. Neutrophils rapidly migrate via lymphatics after Mycobacterium bovis BCG intradermal vaccination and shuttle live bacilli to the draining lymph nodes. Blood 106, 1843–1850 (2005).

30. Rambault, M. et al. Isolation of Bovine Neutrophils by Fluorescence-and Magnetic-Activated Cell Sorting. Methods Mol. Biol. Clifton NJ 2236, 203–217 (2021).

31. Babraham Bioinformatics - FastQC A Quality Control tool for High Throughput Sequence Data. https://www.bioinformatics.babraham.ac.uk/projects/fastqc/.

32. Chen, S., Zhou, Y., Chen, Y. & Gu, J. fastp: an ultra-fast all-in-one FASTQ preprocessor. Bioinformatics 34, i884–i890 (2018).

33. Patro, R., Duggal, G., Love, M. I., Irizarry, R. A. & Kingsford, C. Salmon provides fast and bias-aware quantification of transcript expression. Nat. Methods 14, 417–419 (2017).

34. Soneson, C., Love, M. I. & Robinson, M. D. Differential analyses for RNA-seq: transcript-level estimates improve gene-level inferences. F1000Research 4, 1521 (2015).

35. Love, M. I., Huber, W. & Anders, S. Moderated estimation of fold change and dispersion for RNA-seq data with DESeq2. Genome Biol. 15, 550 (2014).

36. Heavy-tailed prior distributions for sequence count data: removing the noise and preserving large differences | Bioinformatics | Oxford Academic. https://academic.oup.com/bioinformatics/article/35/12/2084/5159452.

37. Benjamini, Y. & Hochberg, Y. Controlling the False Discovery Rate: A Practical and Powerful Approach to Multiple Testing. J. R. Stat. Soc. Ser. B Methodol. 57, 289–300 (1995).

38. Kolberg, L., Raudvere, U., Kuzmin, I., Vilo, J. & Peterson, H. gprofiler2 -- an R package for gene list functional enrichment analysis and namespace conversion toolset g:Profiler. F1000Research 9, ELIXIR-709 (2020).

39. Raudvere, U. et al. g:Profiler: a web server for functional enrichment analysis and conversions of gene lists (2019 update). Nucleic Acids Res. 47, W191–W198 (2019).

40. Ashburner, M. et al. Gene ontology: tool for the unification of biology. The Gene Ontology Consortium. Nat. Genet. 25, 25–29 (2000).

41. Gene Ontology Consortium et al. The Gene Ontology knowledgebase in 2023. Genetics 224, iyad031 (2023).

42. Kobayashi, S. D., DeLeo, F. R. & Quinn, M. T. Microbes and the fate of neutrophils. Immunol. Rev. 314, 210–228 (2023).

43. Morelli, M. P. et al. Neutrophils from tuberculosis patients are polarized toward pro-inflammatory and anti-inflammatory phenotypes according to the disease severity. J. Immunol. 214, 1173–1186 (2025).

44. Cassidy, J. P. et al. Lesions in Cattle Exposed to Mycobacteriumbovis-inoculated Calves. J. Comp. Pathol. 121, 321–337 (1999).

45. Cassidy, J. P. The pathogenesis and pathology of bovine tuberculosis with insights from studies of tuberculosis in humans and laboratory animal models. Vet. Microbiol. 112, 151–161 (2006).

46. Wang, J. et al. Expression pattern of interferon-inducible transcriptional genes in neutrophils during bovine tuberculosis infection. DNA Cell Biol. 32, 480–486 (2013).

47. Wang, J. et al. Investigation of the effect of Mycobacterium bovis infection on bovine neutrophils functions. Tuberc. Edinb. Scotl. 93, 675–687 (2013).

48. Shu, D. et al. Comparison of gene expression of immune mediators in lung and pulmonary lymph node granulomas from cattle experimentally infected with *Mycobacterium bovis*. Vet. Immunol. Immunopathol. 160, 81–89 (2014).

49. Dorhoi, A. et al. Type I IFN signaling triggers immunopathology in tuberculosis_susceptible mice by modulating lung phagocyte dynamics. Eur. J. Immunol. 44, 2380–2393 (2014).

50. Gern, B. H. et al. Early and opposing neutrophil and CD4 T cell responses shape pulmonary tuberculosis pathology. J. Exp. Med. 222, e20250161 (2025).

51. Branchett, W. J. et al. Type I IFN drives neutrophil swarming, impeding lung T cell–macrophage interactions and TB control. J. Exp. Med. 222, e20250466 (2025).

52. Maertzdorf, J. et al. Functional Correlations of Pathogenesis-Driven Gene Expression Signatures in Tuberculosis. PLoS ONE 6, e26938 (2011).

53. Remot, A. et al. Mycobacterial Infection of Precision-Cut Lung Slices Reveals Type 1 Interferon Pathway Is Locally Induced by Mycobacterium bovis but Not M. tuberculosis in a Cattle Breed. Front. Vet. Sci. 8, 696525 (2021).

54. Chunfa, L. et al. The Central Role of IFI204 in IFN-β Release and Autophagy Activation during Mycobacterium bovis Infection. Front. Cell. Infect. Microbiol. 7, 169 (2017).

55. Wang, J. et al. Inhibition of type I interferon signaling abrogates early Mycobacterium bovis infection. BMC Infect. Dis. 19, 1031 (2019).

56. Pedrosa, J. et al. Neutrophils Play a Protective Nonphagocytic Role in Systemic Mycobacterium tuberculosis Infection of Mice. Infect. Immun. 68, 577–583 (2000).

57. Corleis, B. et al. Escape of Mycobacterium tuberculosis from oxidative killing by neutrophils. Cell. Microbiol. 14, 1109–1121 (2012).

58. Jones, G. S., Amirault, H. J. & Andersen, B. R. Killing of Mycobacterium tuberculosis by neutrophils: a nonoxidative process. J. Infect. Dis. 162, 700–704 (1990).

59. Majeed, M., Perskvist, N., Ernst, J. D., Orselius, K. & Stendahl, O. Roles of calcium and annexins in phagocytosis and elimination of an attenuated strain of Mycobacterium tuberculosis in human neutrophils. Microb. Pathog. 24, 309–320 (1998).

60. Knaul, J. K. et al. Lung-Residing Myeloid-derived Suppressors Display Dual Functionality in Murine Pulmonary Tuberculosis. Am. J. Respir. Crit. Care Med. 190, 1053–1066 (2014).

61. Agrawal, N. et al. Human Monocytic Suppressive Cells Promote Replication of Mycobacterium tuberculosis and Alter Stability of in vitro Generated Granulomas. Front. Immunol. 9, (2018).

62. Tsiganov, E. N. et al. Gr-1dimCD11b+ immature myeloid-derived suppressor cells but not neutrophils are markers of lethal tuberculosis infection in mice. J. Immunol. 192, 4718–4727 (2014).

63. Doz-Deblauwe, É. et al. CR3 Engaged by PGL-I Triggers Syk-Calcineurin-NFATc to Rewire the Innate Immune Response in Leprosy. Front. Immunol. 10, (2019).

64. Voskuil, M. I., Bartek, I. L., Visconti, K. & Schoolnik, G. K. The Response of Mycobacterium Tuberculosis to Reactive Oxygen and Nitrogen Species. Front. Microbiol. 2, (2011).

65. Vilchèze, C., Hartman, T., Weinrick, B. & Jacobs, W. R. Mycobacterium tuberculosis is extraordinarily sensitive to killing by a vitamin C-induced Fenton reaction. Nat. Commun. 4, 1881 (2013).

66. Fan, X.-Y. et al. Oxidation of dCTP contributes to antibiotic lethality in stationary-phase mycobacteria. Proc. Natl. Acad. Sci. 115, 2210–2215 (2018).

67. Akhtar, P. et al. Rv3303c of Mycobacterium tuberculosis protects tubercle bacilli against oxidative stress in vivo and contributes to virulence in mice. Microbes Infect. 8, 2855–2862 (2006).

68. Schmielau, J. & Finn, O. J. Activated granulocytes and granulocyte-derived hydrogen peroxide are the underlying mechanism of suppression of t-cell function in advanced cancer patients. Cancer Res. 61, 4756–4760 (2001).

69. Kusmartsev, S., Nefedova, Y., Yoder, D. & Gabrilovich, D. I. Antigen-Specific Inhibition of CD8+ T Cell Response by Immature Myeloid Cells in Cancer Is Mediated by Reactive Oxygen Species. J. Immunol. 172, 989–999 (2004).

70. Frossi, B., De Carli, M., Piemonte, M. & Pucillo, C. Oxidative microenvironment exerts an opposite regulatory effect on cytokine production by Th1 and Th2 cells. Mol. Immunol. 45, 58–64 (2008).

71. Bassoy, E. Y., Walch, M. & Martinvalet, D. Reactive Oxygen Species: Do They Play a Role in Adaptive Immunity? Front. Immunol. 12, 755856 (2021).

72. Corzo, C. A. et al. Mechanism Regulating Reactive Oxygen Species in Tumor-Induced Myeloid-Derived Suppressor Cells. J. Immunol. 182, 5693–5701 (2009).

73. Rice, C. M. et al. Tumour-elicited neutrophils engage mitochondrial metabolism to circumvent nutrient limitations and maintain immune suppression. Nat. Commun. 9, 5099 (2018).

74. Geering, B. & Simon, H.-U. Peculiarities of cell death mechanisms in neutrophils. Cell Death Differ. 18, 1457–1469 (2011).

75. Chen, T., Ren, Q. & Ma, F. New insights into constitutive neutrophil death. Cell Death Discov. 11, 6 (2025).

76. Schaaf, K. et al. Mycobacterium tuberculosis exploits the PPM1A signaling pathway to block host macrophage apoptosis. Sci. Rep. 7, 42101 (2017).

77. Behar, S. M. et al. Apoptosis is an innate defense function of macrophages against Mycobacterium tuberculosis. Mucosal Immunol. 4, 279–287 (2011).

78. Pfrommer, E. et al. Enhanced tenacity of mycobacterial aerosols from necrotic neutrophils. Sci. Rep. 10, 9159 (2020).

79. van der Meer, A. J. et al. Neutrophil extracellular traps in patients with pulmonary tuberculosis. Respir. Res. 18, 181 (2017).

80. Moreira-Teixeira, L. et al. Type I IFN exacerbates disease in tuberculosis-susceptible mice by inducing neutrophil-mediated lung inflammation and NETosis. Nat. Commun. 11, 5566 (2020).

81. Ramos-Kichik, V. et al. Neutrophil extracellular traps are induced by *Mycobacterium tuberculosis*. Tuberculosis 89, 29–37 (2009).

82. Filio-Rodríguez, G. et al. In vivo induction of neutrophil extracellular traps by Mycobacterium tuberculosis in a guinea pig model. Innate Immun. 23, 625–637 (2017).

83. Rustom, A., Saffrich, R., Markovic, I., Walther, P. & Gerdes, H.-H. Nanotubular Highways for Intercellular Organelle Transport. Science 303, 1007–1010 (2004).

84. Ariazi, J. et al. Tunneling Nanotubes and Gap Junctions–Their Role in Long-Range Intercellular Communication during Development, Health, and Disease Conditions. Front. Mol. Neurosci. 10, 333 (2017).

85. André, A. C. et al. Proteoglycofili are glycosaminoglycan-containing fibrillar components released by neutrophils to neutralize bacteria. Cell Rep. 44, 116541 (2025).

86. Paape, et al. The bovine neutrophil: Structure and function in blood and milk. Vet. Res. 34, 597–627 (2003).

87. Tan, B. H. et al. Macrophages Acquire Neutrophil Granules for Antimicrobial Activity against Intracellular Pathogens. J. Immunol. 177, 1864–1871 (2006).

88. Dallenga, T. et al. M. tuberculosis-Induced Necrosis of Infected Neutrophils Promotes Bacterial Growth Following Phagocytosis by Macrophages. Cell Host Microbe 22, 519–530.e3 (2017).

