## Supplementary material for "Functional divergence of regulatory and conventional bovine neutrophils following *Mycobacterium bovis* infection": SupFig1

### Supplementary Figure 1

**a**

GeneOntology Biological Processes
BCG-infected $N^{reg}$ *versus* $N^{conv}$

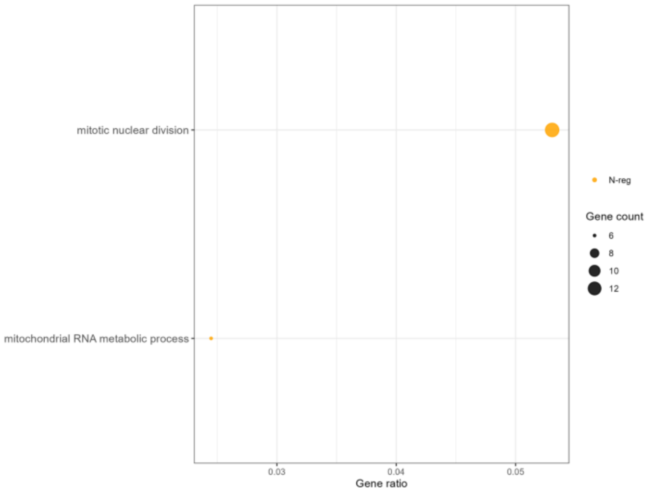


**b**

GeneOntology Biological Processes
Mb3601-infected $N^{reg}$ *versus* $N^{conv}$

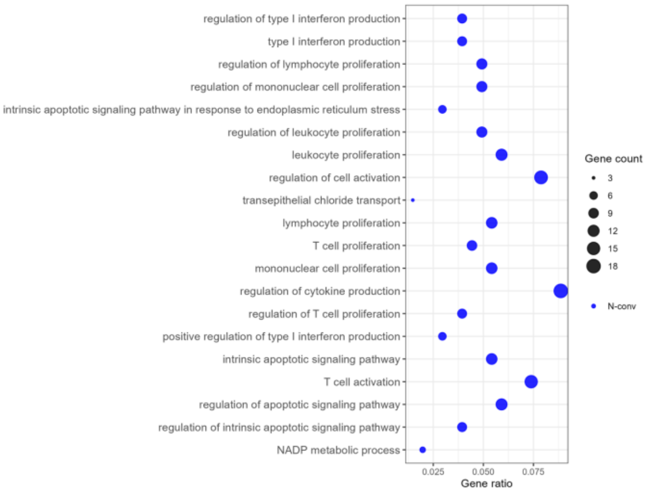
