## Supplementary figures and images for "Functional divergence of regulatory and conventional bovine neutrophils following *Mycobacterium bovis* infection"

### SupFig2

#### Supplementary Figure 2

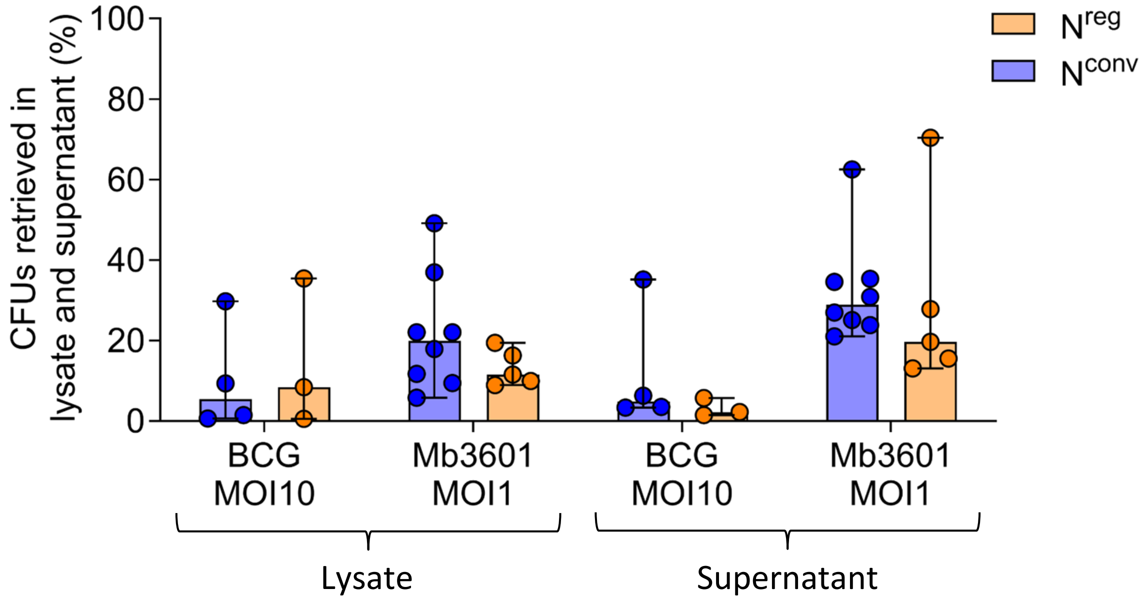
